# Systematic assessment of transcriptomic and phenotypic biological profiling for mechanism-based hazard assessment using target-annotated reference chemicals in renal proximal tubular epithelial cells

**DOI:** 10.64898/2026.08.08.743660

**Authors:** Hugo W. van Kessel, Marlene Wedler, Palle Helmke, Dace Žīgure, Stephen S. Ferguson, Joshua Harrill, Gerhard Ecker, Shu Liu, Michael Oelgeschläger, Giulia Callegaro, Bob van de Water

## Abstract

Integrating high-throughput in vitro data into next-generation risk assessment (NGRA) workflows requires screening strategies that yield quantitative potency estimates and mechanistically interpretable biological signals. Transcriptomic and morphological profiling are increasingly adopted for early-stage hazard identification by enabling triage of substances for resource-intensive follow-up and prioritizing candidates most likely to present meaningful risk. In this study, we aimed to characterize biological concordance and uncertainty by quantifying how well high-throughput transcriptomics (HTTr) and Cell Painting PLUS (CPP) bioactivity profiles recover target-relevant biological signals in immortalized human renal proximal tubule epithelial RPTEC/TERT1 cells using 313 reference chemicals with high-confidence target annotations. Through quality control procedures and biological activity filters we yielded 142 reference chemicals spanning 66 different targets, which were systematically evaluated for biological concentration-responses by HTTr and CPP. HTTr was evaluated using TXG-MAPr-based qualitative and quantitative gene network activity analysis. HTTr showed the most prominent activity for targets that were highest expressed in RPTEC/TERT1 cells. Active chemical-pairs showed strong gene network activity correlation albeit with different potencies. Similarly, the highest transcriptomic concordance was observed for reference chemicals acting in the same pathway, such as EGFR/MEK or PI3K/AKT/mTOR. CPP often showed high sensitivity primarily at the organelle level providing limited statistical power for chemical grouping. Collectively, the results support HTTr and CPP as complementary early-tier assays within an in vitro weight-of-evidence safety testing framework. Although CPP is suitable as a cost-effective screening modality, HTTr offers higher mechanistic resolution for mode-of-action inference in high-throughput bioactivity screening and therefore remains necessary for high-confidence mechanistic interpretation.

## 1 Introduction

Regulators, industry, and academia increasingly converge on next-generation risk assessment (NGRA) frameworks that incorporate new approach methodologies (NAMs) to improve human relevance, accelerate decision-making, and reduce reliance on animal testing (Cronin et al., 2025; Moné et al., 2020; Pallocca et al., 2022; Thomas et al., 2019). Within this context, early-stage hazard identification is positioned as a critical gatekeeping step that prioritizes substances for deeper evaluation and focuses downstream resources on the most plausible risks (Leist et al., 2025).

Two high-throughput screening modalities are increasingly adopted to support early hazard characterization. Firstly, high-throughput transcriptomic (HTTr) assay allows the cost-effective characterization of concentration- and time-dependent transcriptional responses after chemical exposure (Ye et al., 2018; Yeakley et al., 2017). Toxicogenomic interpretation frameworks, such as TXG-MAPr, support the qualitative and quantitative analysis of HTTr data and enable the mechanistic interpretation of transcriptional perturbations within a test system-specific context, thereby facilitating the biological grouping of substances (Callegaro et al., 2023; Kunnen et al., 2025; van Kessel et al., 2026; Vrijenhoek et al., 2022; Wijaya et al., 2024). Early transcriptional changes related to adverse outcomes may then initiate further testing in more complex (multi-cellular) test systems that reflect human (patho)physiology. Secondly, Cell Painting (CP) is a high-content morphological profiling approach that captures phenotypic perturbations across cellular compartments (States et al., 2013). Recently, the Cell Painting PLUS (CPP) approach has been established that includes phenotypic assessment of additional organelles (von Coburg et al., 2025). Although the underlying mechanisms of the phenotypic changes are typically unknown, the phenotypic changes reflect chemical-biological interactions that could be reflective for toxicological perturbations. While HTTr and CP(P) can capture hazards in an early tier testing, an important remaining question is how well bioactivity profiles are recovered by HTTr and CP(P) across a diverse panel of pharmacological and toxicological relevant targets and how consistent these biological responses are for substances that have a similar target and/or mode of action.

To translate screening readouts into actionable metrics for risk assessment, quantitative methods such as benchmark concentration (BMC) modeling are applied to derive in vitro points of departure (PODs) (Harrill et al., 2021; Johnson et al., 2025; Judson et al., 2024). Short term transcriptomic PODs (tPODs) correlate with apical chronic PODs derived from in vivo studies, highlighting their relevance and use as predictive indicators of toxicity (Farmahin et al., 2017; Pagé-Larivière et al., 2019). The TXG-MAPr toxicogenomic interpretation framework supports test system specific tPOD determination of co-expression gene networks, thus providing mechanistic context for tPODs. CP(P) data have been analyzed using two complementary approaches to derive PODs (Nyffeler et al., 2020; Way et al., 2022). First, BMCs were calculated for each feature using BMC modeling to compare and cluster compounds based on the fraction of significantly perturbed features within each feature category. Second, phenotype altering concentrations (PACs), corresponding to CPP PODs, were derived from all features for each compound for comparison with tPODs. Recently, several studies have evaluated the concordance and complementarity of HTTr and High-throughput phenotypic profiling (HTPP) for chemical bioactivity assessment (Harrill et al., 2024a; Jurgelewicz et al., 2025). A systematic assessment of the concordance between HTTr and CPP of substances with similar pharmacological and/or toxicological molecular targets is so far lacking.

Here, we evaluated HTTr and CPP performance in RPTEC/TERT1 cells using 313 reference chemicals with high-confidence molecular target annotations, including at least two chemicals per target (Judson et al., 2018a; Wieser et al., 2008). We aimed to quantify the extent to which transcriptomic and morphological bioactivity profiles capture target-relevant biological signals, support mechanistic inference, and enable potency estimation under harmonized experimental conditions. In short, reference chemicals verified by mass spectrometry were evaluated for transcriptomic activity after 24 hours of exposure at the highest non-cytotoxic concentration, and chemical-target groups were advanced for further testing if at least one chemical elicited activity. A set of 142 reference chemicals spanning 66 cellular targets was evaluated using a seven-point concentration-response design in HTTr and CPP at 24 hours. Using HTTr, we characterized downstream cellular responses following target engagement by reference chemicals and linked the resulting responses to the intended molecular targets. In summary, our findings support the use of HTTr and CPP as early-stage modalities within an in vitro weight-of-evidence nephrotoxicity assessment pipeline. Our results support HTTr in combination with TXG-MAPr analysis as the most informative method to provide data-driven mechanistic inference of toxicological hazard.

## 2 Materials and methods

### 2.1 RPTEC/TERT1 cell culture

RPTEC/TERT1 cells were obtained under license from Evercyte GmBH (Vienna, Austria) and cultured between passages 40 and 45. Unless stated otherwise, cells were cultured in 96-well ScreenStar plates (Greiner, 655866) pre-coated with 0.1 mg/mL Collagen I (Sigma-Aldrich, C3867), at a seeding density of 60,000 cells/cm² and maintained at 37 °C in a humidified atmosphere containing 5% CO₂. Following confluency, cells were cultured for an additional 10 days. The culture medium consisted of a 1:1 mixture of Ham’s F-12 Nutrient Mix (Thermo Fisher Scientific, 21765029) and Dulbecco’s Modified Eagle Medium without glucose (Thermo Fisher Scientific, 11966025), supplemented with 50 U/mL penicillin and 50 µg/mL streptomycin (Thermo Fisher Scientific, 15070063), 5 µg/mL insulin, 5 µg/mL transferrin, and 5 ng/mL sodium selenite (Sigma-Aldrich, I1884) reconstituted sequentially in 10 mM glacial acetic acid (Biosolve, 0001080501BS) and PBS (Sigma-Aldrich, D8537). Additional supplements included 2 mM GlutaMAX (Thermo Fisher Scientific, 35050061), 36 ng/mL hydrocortisone (Sigma-Aldrich, H0135) reconstituted in ethanol (Sigma-Aldrich, 29221) and PBS, and 10 ng/mL human epidermal growth factor (Sigma-Aldrich, E9644) reconstituted in 10 mM glacial acetic acid containing 0.1% (w/v) BSA (Sigma-Aldrich, A9647). The medium was further supplemented with 0.5% (v/v) fetal bovine serum (TICO Europe, FBSEU500) and changed every two days.

### 2.2 Reference chemicals

#### 2.2.1 Library assembly and target annotation

A collection of 313 reference chemicals with known cellular targets was assembled using RefChemDB, a resource containing chemical-target annotations generated through a semi-automated curation process based on public databases such as ToxCast and ChEMBL (Judson et al., 2018a). Each annotation included a chemical identifier, the corresponding cellular target, and the type of modulation. To ensure reliability, chemicals were filtered to retain only those with a support level ≥ 5, reflecting evidence from at least five distinct literature sources (Bundy et al., 2024). For each selected target, at least two associated chemicals were obtained, allowing for comparative analysis of chemicals acting on the same target. Chemicals were obtained from the U.S. EPA’s chemical contractor EvoTec (Princeton, NJ) and provided in LabCyte Echo-qualified 384-well low dead volume (384LDV) plates as 20 mM stock solutions or the highest soluble concentration in DMSO and stored at −80 °C until use.

#### 2.2.2 Practical implications

In RefChemDB, individual chemicals may be linked to multiple cellular targets, resulting in non–mutually exclusive associations. To reduce ambiguity, each chemical was assigned to a single target by retaining the linkage with the highest support level. The selection of reference chemicals was based solely on the availability of high-confidence target information, without consideration of potency, efficacy, or pharmacokinetic properties, and was therefore cell type–agnostic. Target annotations were represented by gene symbols corresponding to their encoded protein products. These annotations should be interpreted as functional target proxies rather than strictly unique molecular targets. In some cases, these symbols denote higher-order protein assemblies or homologous protein groups rather than discrete individual proteins, as exemplified by TUBA1A for tubulin. Accordingly, the retained target should be interpreted as the best-supported representative annotation, rather than evidence that related proteins or protein complexes are not relevant targets. Chemical identities were represented using the preferred name whenever available. When unavailable, the U.S. EPA DSSTox Substance Identifier (DTXSID) was used.

#### 2.2.3 Cross-database mapping for bioactivity and target annotation

To characterize the bioactivity of reference chemicals, pChEMBL values were obtained from ChEMBL release 33 (Mendez et al., 2019). Briefly, reference chemicals were mapped using DTXSID identifiers, which were uploaded to the EPA CompTox Chemicals Dashboard (Williams et al., 2017) to retrieve standardized chemical descriptors. For each reference chemical, all ChEMBL molecular forms linked to the primary InChIKey were retrieved, and human bioactivity records for these forms were queried via the ChEMBL API to extract the corresponding pChEMBL values. In addition, reference chemical targets were mapped to ChEMBL target identifiers and cross-referenced to UniProt identifiers (Bateman et al., 2025), which were queried via the UniProt API to retrieve protein annotations including subcellular location and cellular function. As the obtained target annotations were heterogeneous and frequently comprised multiple annotations per target, they were manually curated to reduce redundancy and ambiguity to improve their suitability for visualization.

### 2.3 Mass spectrometry

#### 2.3.1 Liquid chromatography–high-resolution mass spectrometry (LC-HRMS)

Chromatographic analyses were performed using an Acquity UPLC H-Class system (Waters, Milford, MA, USA) coupled to a Synapt G2-Si high-resolution time-of-flight (TOF) mass spectrometer (Waters). Separation was achieved on an Acquity BEH C18 column (2.1 × 50 mm, 1.7 μm) maintained at 40 °C. The mobile phase consisted of 0.1% (v/v) formic acid in water and acetonitrile. The following gradient was applied: initial 95% aqueous phase and 5% acetonitrile, decreased linearly to 2% aqueous phase and 98% acetonitrile at 1.5 minutes, maintained at this composition until 2.5 minutes, returned to 95% aqueous phase and 5% acetonitrile at 3 minutes, and held at this composition until 4 minutes. The flow rate was 0.4 mL/minutes, and the injection volume was 1 μL. Mass spectrometric detection was performed using electrospray ionization (ESI) in both positive and negative ion modes. In ESI positive mode, the capillary voltage was set at 0.7 kV and the cone voltage at 40 V. In ESI negative mode, the capillary voltage was 2.5 kV and the cone voltage 40 V. The source temperature was 120 °C, the desolvation gas (nitrogen) flow rate was 800 L/hour, and the desolvation temperature was 400 °C. Leucine enkephalin was used as the lock mass for mass calibration. The TOF mass analyzer was operated in full-scan mode across the m/z range of 50–1200. For selected samples, direct infusion analysis was performed without chromatographic separation.

#### 2.3.2 Gas chromatography–mass spectrometry (GC-MS)

GC-MS analyses were conducted using an Agilent 7890A gas chromatograph coupled to an Agilent 5975C mass selective detector (Agilent Technologies, Santa Clara, CA, USA). An Agilent HP-5MS capillary column (30 m length, 0.25 mm inner diameter, 0.25 μm film thickness) was used, with helium as the carrier gas at a constant flow rate of 0.927 mL/minutes. The injector temperature was 250 °C, operating in splitless mode, with an injection volume of 1 μL and a solvent delay of 3.6 minutes. The oven temperature program was as follows: initial temperature of 80 °C held for 2 minutes, increased at 10 °C/minute to 260 °C, and held for 20 minutes, resulting in a total run time of 40 minutes. Mass spectrometric detection was performed in electron ionization (EI) positive mode.

A total of 313 reference chemicals were analyzed using the described LC-HRMS and GC-MS methods. The combined use of both techniques ensured detection coverage across analytes with diverse physicochemical properties. Twenty-six compounds were not correctly detected under the applied conditions and were consequently excluded from further analyses and experimental evaluation (Tab. S1).

### 2.4 Cell viability assessment

RPTEC/TERT1 cells were cultured in 384-well ScreenStar plates (Greiner Bio-One, 781866) and reference chemicals were tested at 24 hours using eight concentrations spaced at half-log10 intervals decreasing from a maximum of 30 µM, with 0.15% (v/v) DMSO maintained across all conditions. For compounds with limited solubility, the maximum concentration was the highest soluble concentration achievable at 0.15% (v/v) DMSO. Each condition was performed in four biological replicates in a total well volume of 100 µL. Negative controls containing 0.15% DMSO (v/v) were included on every plate. Conditions for a given reference chemical were not distributed across different plates within a biological replicate to minimize plate variability. Two hours before exposure, cells were stained with 100 ng/mL Hoechst 33342 and then replaced with reference chemicals in medium containing 0.2 µM propidium iodide (PI). Image acquisition was performed using a Nikon TiE2000 laser confocal microscope inside a 37 °C, 5% CO₂ incubator, equipped with an automated stage, perfect focus, and 405/561 nm lasers to detect Hoechst and PI signals. Raw images were extracted with Nikon NIS Elements Viewer v5.21, and nuclei were segmented using watershed masked clustering (Di et al., 2012) in Fiji (Schindelin et al., 2012). Binary Hoechst and raw PI images were analyzed in CellProfiler v4.2.5 (Stirling et al., 2021) using modules to identify, relate, and quantitatively measure objects. Nuclei were classified as PI-positive when PI objects overlapped more than one third of the nucleus area. Reference chemicals were classified as cytotoxic when the fraction of PI-positive cells exceeded 30% under a given condition. The concentration range was adjusted for these chemicals to enable subsequent analyses (Tab. S1).

### 2.5 HTTr data generation

#### 2.5.1 HTTr activity evaluation

High-throughput transcriptomic profiling was conducted using the TempO-Seq platform (BioClavis Ltd. Glasgow, UK) in two sequential stages (Yeakley et al., 2017). The first stage involved a transcriptional activity evaluation screen, designed to determine whether exposure to a high concentration of each test chemical elicited a measurable transcriptional response after 24 hours. Concentrations were set at 30 µM wherever feasible; however, for compounds with limited solubility in DMSO, lower maximum concentrations were applied (Tab. S1). Following 24 hours of exposure, cells were washed with phosphate-buffered saline (PBS) and lysed in 50 µL of a 1:1 (v/v) solution of 2× TempO-Seq lysis buffer and PBS for 15 minutes at room temperature. Lysates were snap-frozen on dry ice, stored at –80 °C, and subsequently shipped on dry ice for processing using the TempO-Seq human whole-transcriptome probe panel version 2.0, comprising 22537 probes targeting 19703 genes. Differential gene expression profiles were generated as described in Section 2.6. Chemicals groups associated with the same molecular target were selected for further analysis if at least one compound in the group induced twenty or more differentially expressed genes, defined by an absolute log2FC greater than 1.5 and an adjusted p-value below 0.05.

#### 2.5.2 HTTr concentration–response profiling

The second stage consisted of a concentration–response profiling screen to quantify transcriptional activity with the objective of determining transcriptional points of departure (tPODs). This phase targeted 66 cellular targets using a panel of 142 chemicals, each selected based on transcriptomic activity observed in the transcriptional activity evaluation screen. As positive controls, a seven-point concentration range of cisplatin and CDDO-Me was included, using two-fold dilution steps starting at 100 µM and 1 µM, respectively. All other reference chemicals were tested in a seven-point concentration series with half-log10 spacing. The final DMSO concentration was maintained at 0.15% (v/v) across all conditions. Each 96-well assay plate included ten control samples: three medium-only, three vehicle control of 0.15% (v/v) DMSO, three treated with 100 µM cisplatin as a positive control, and MAQC which was added post-exposure on the shipment plates. Cell lysis and sample preparation followed the same procedure as described in Section 2.5.1. Samples were processed using the human whole-transcriptome TempO-Seq probe panel version 2.1, comprising 22533 probes targeting 19683 genes. The concentration–response profiling screen was conducted in three biological replicates, each consisting of 25 assay plates, resulting in a total of 3942 samples. Quality control metrics for the concentration–response profiling screen are presented in Figure S1.

### 2.6 HTTr DGE analysis

The vendor performed quality control and alignment of the FASTQ files and provided a count matrix with probes in rows and samples in columns. The count matrix was pre-processed to remove low-expression probes and low-quality samples prior to differential gene expression (DGE) analysis. First, probes without an Entrez Gene ID in the vendor-provided manifest and MAQC control samples, which are not relevant to the DGE analysis, were excluded. The remaining data were normalized using the counts per million (CPM) method, followed by probe-level filtering using the relevance filter from the R-ODAF framework to remove low-expression probes (Verheijen et al., 2022). The original count matrix was then filtered to retain only the probes that passed these filtering steps. Before sample-level filtering, and after excluding MAQC controls, the median number of mapped reads across 3867 samples was 2.73 million, with a mean of 2.78 million and an interquartile range of 2.10 to 3.39 million mapped reads. Next, sample-level filtering was applied by removing samples with fewer than 0.9 million mapped reads. Probe counts were then aggregated to the gene level by summing counts for probes mapping to the same gene. Gene-level counts were then CPM normalized and log2(X + 1) transformed (Everett et al., 2022). To assess replicate consistency, Pearson correlations were calculated between each replicate and a pseudo-sample, defined as the mean expression profile across all replicates within a condition. Replicates with a correlation below 0.8 were excluded from further analysis. The original count matrix was subsequently filtered to retain only the probes and samples that passed all prior filtering steps. Following this filtering, DGE analysis was performed using DESeq2 v1.46 (Love et al., 2014) in R v4.4.2 (R Core Team, 2025). For each chemical, samples across all concentrations and time points were jointly modeled with plate-matched vehicle controls to extract relevant contrasts and generate DGE profiles.

### 2.7 HTTr BMC modeling and functional interpretation

Two methods for module tPOD determination were performed: one based on log2 CPM-normalized counts and the other on module eigengenes (MEs). For both approaches, raw read counts underwent probe- and sample-level pre-processing as described in Section 2.6. In the log2(CPM)-based method, the pre-processed count matrix was normalized to counts per million (CPM), followed by a log2(X + 1) transformation. In the ME-based method, differential gene expression analysis was performed using DESeq2 v1.46 in R v4.4.2 to generate replicate-specific differential gene expression profiles by comparing individual replicates to plate-matched controls (Kunnen et al., 2024). These replicate-level DGE profiles were uploaded to the RPTEC/TERT1 TXG-MAPr platform to compute ME profiles for each replicate. Subsequently, for each method, the resulting data were organized into separate tables per reference chemical. Each table contained the biological replicates for all tested concentrations along with their corresponding plate-matched controls. These controls included both technical and biological replicates from the same experimental plates and served as the biological baseline.

In BMDExpress 3 v3.20 (Phillips et al., 2018; Yang et al., 2007), we performed William’s Trend Test (WTT) with p < 0.05 and 10.000 permutations. In the log2(CPM)-based method, WTT included a fold-change threshold greater than 1.5. Benchmark concentration–response modeling was performed using a benchmark response (BMR) of 1.349 standard deviations (10%) (O’Brien et al., 2025). BMC fits were retained if they met the following quality criteria: a BMC/BMCL ratio below 20, a BMCU/BMC ratio below 20, a coefficient of determination (R2) of at least 0.6, and a BMC value within the range of 10% of the lowest tested concentration to the highest tested concentration. In the log2(CPM)-based method, for each reference chemical, the modules from the RPTEC/TERT1 TXG-MAPr were used as gene sets, and tPODs were calculated as the median gene BMC for each module, provided there was an overlap of at least three genes and coverage of at least 5% of the module and at least one gene in the module was identified as a DEG during HTTr data analysis. In the ME-based method, module BMCs were directly used as tPODs, retaining only modules with an absolute ME > 1 in at least one condition for each reference chemical. Functional interpretation of modules was supported by knowledge-based pathway enrichment and regulatory inference tools integrated into the human in vitro RPTEC/TERT1 TXG-MAPr (van Kessel et al., 2026). All reported correlation coefficients were computed using Pearson’s correlation.

### 2.8 CPP data generation

RPTEC/TERT1 cells were cultured in 384-well ScreenStar plates (Greiner Bio-One, 781866) under the conditions described in Section 2.1. The chemical set comprised the same 142 reference chemicals selected for the HTTr concentration–response profiling screen described in Section 2.5.2. After 24 hours of chemical exposure, staining and imaging were conducted as described by in five biological replicates. Image acquisition was performed using an Opera Phenix High-Content Screening System (Revvity Inc.) in confocal mode with a 20x water objective with three fields captured per well. Four sequential imaging channels were used, each defined by the excitation laser wavelength followed by the corresponding emission detection range: 405/435–480, 488/500–550, 561/570–630, and 640/650–760 nm. During the first staining cycle, cells were incubated with Cell Navigator™ Lysosome Staining Kit NIR Fluorescence with LysoBrite™ NIR lysotropic dye (AAT Bioquest, 22652) in a 1:500 dilution and 0.5 µM MitoTracker Orange CMTMROS (Thermo Fisher Scientific, M7510) for 30 min at 37 °C, and subsequently fixed in 4% PFA for 20 min. After three PBS washes, cells were incubated for 30 minutes with 0.1% Triton X-100, 33 nM Alexa Fluor™ Plus 405 Phalloidin (Thermo Fisher Scientific, A30104), and 0.3 µM SYTO™ 14 Green Fluorescent Nucleic Acid Stain (Thermo Fisher Scientific, S7576). The first staining cycle was completed with three PBS washes followed by image acquisition. The first cycle was succeeded by dye elution, in which all dyes except MitoTracker were eluted through three washes with deionized water followed by a 10-minute incubation in elution buffer containing 0.5 M L-glycine and 1% SDS (pH 2.5) and washed three times with water and three times with PBS. During the second staining cycle, cells were incubated for 30 minutes with 4 µM Hoechst 33342 (Thermo Fisher Scientific, H3570), 1.3 µg/ml Wheat Germ Agglutinin Alexa Fluor 488 Conjugate (Thermo Fisher Scientific, W11261) and 10 µg/ml Concanavalin A-Alexa Fluor 647 Conjugate (Thermo Fisher Scientific, C21421). The second staining cycle was completed with three PBS washes followed by image acquisition.

### 2.9 CPP concentration-response modeling

Image analysis and BMC modelling were performed as previously (von Coburg et al., 2025) described using Harmony v4.8 (Revvity Inc.) and tcplfit2 (Sheffield et al., 2022) v0.1.9 in R v4.4.2, respectively. Initially, images from both staining cycles were merged into a single multi-stack image using the application specific Add Channels 4i building block (Kirsch, 2025). Subsequently, cell segmentation and feature extraction were carried out using a customized analysis pipeline. Single-cell data were aggregated to the well level by calculating the median value for each feature across all cells within a well. The well data were normalized using B-score normalization, which computes a robust z-score after removing row and column effect using median-based statistics (Mpindi et al., 2015) to obtain a B-score and correct the plate-level patterns as described in <u>Wedler et al. 2026 (in preparation)</u>. For each compound and feature combination, concentration–response modelling was performed using five models (constant, hill, poly1, poly2, and power), with the BMR defined as 1.349 standard deviations (10%). BMC values were classified as invalid when the hit-call was below 0.9, when either the BMCU or BMCL was missing, or when the BMC exceeded the highest tested concentration or fell below one tenth of the lowest tested concentration. Each feature was annotated according to imaging channel (e.g., DNA, ER, RNA), cellular region (e.g., cytoplasm, nucleus, membrane), and type (e.g., texture, intensity, compactness). The combination of these attributes defined the feature category, e.g., DNA–nucleus–compactness. To determine the phenotype-altering concentration (PAC) for each chemical, 894 features were summarized using Mahalanobis distance–based fitting (Nyffeler et al., 2020). Briefly, all individual features and composite feature categories were reduced in dimensionality using principal component analysis (PCA). Mahalanobis distances between treatment and control groups were then computed from the PCA-transformed feature set and modeled using tcplfit2, with the BMR defined as 2MAD. The lowest BMC among the global BMCs derived from all features and those obtained for each feature category was defined as the PAC. In addition, for each chemical, a one-sided t-test was performed for every feature to assess whether the feature value for a chemical was significantly lower (p<0.05) than the corresponding feature values across all other chemicals. For each channel–region feature category (e.g. DNA–nucleus), the fraction of significant features was defined as the number of significant features divided by the total number of features. A chemical was classified as active if at least one category exhibited a fraction of significant BMCs greater than 30%. Notably, the generic membrane feature category exhibited a binary response, with compounds either active or inactive.

## 3 Results

We defined an experimental workflow outlining the sequential data generation and processing steps used to establish the HTTr and CPP datasets (Fig. 1A). To better characterize the biological target space, cellular function and subcellular localization of the targets of our 313 substances, data were extracted from UniProt. The targets exhibited substantial diversity, with a large proportion corresponding to membrane-associated receptors followed by intracellular transferases and oxidoreductases (Fig. 1B; Table S1). Mass spectrometry verified reference chemicals were evaluated for transcriptomic activity at the highest non-cytotoxic concentration following 24-hour exposure. Chemical-target groups were retained when at least one chemical exhibited transcriptional activity, defined by a minimum of 20 differentially expressed genes (DEGs). This selection yielded 142 chemicals, which were subsequently tested in both HTTr and CPP assays in a concentration-response setting across seven concentrations at a single 24-hour time point. BMC analysis was conducted separately for each assay to derive tPODs and CPP-derived PACs. Module tPODs were derived using two complementary representations. In the log2(CPM)-derived gene-level approach, module tPODs were calculated as the median gene BMC within each module. In the module eigengene (ME) approach, MEs were computed from replicate-level DESeq2 differential expression profiles and module BMCs were directly used as tPODs. Both approaches were applied because the relative suitability of gene-level aggregation versus direct module-level modeling for module tPOD estimation is not established. Nonetheless, among modules for which tPODs were obtained with both methods, log2(CPM)-derived and ME-derived module tPODs were strongly correlated (Pearson r = 0.82; Fig. S2).

**Figure 1.**
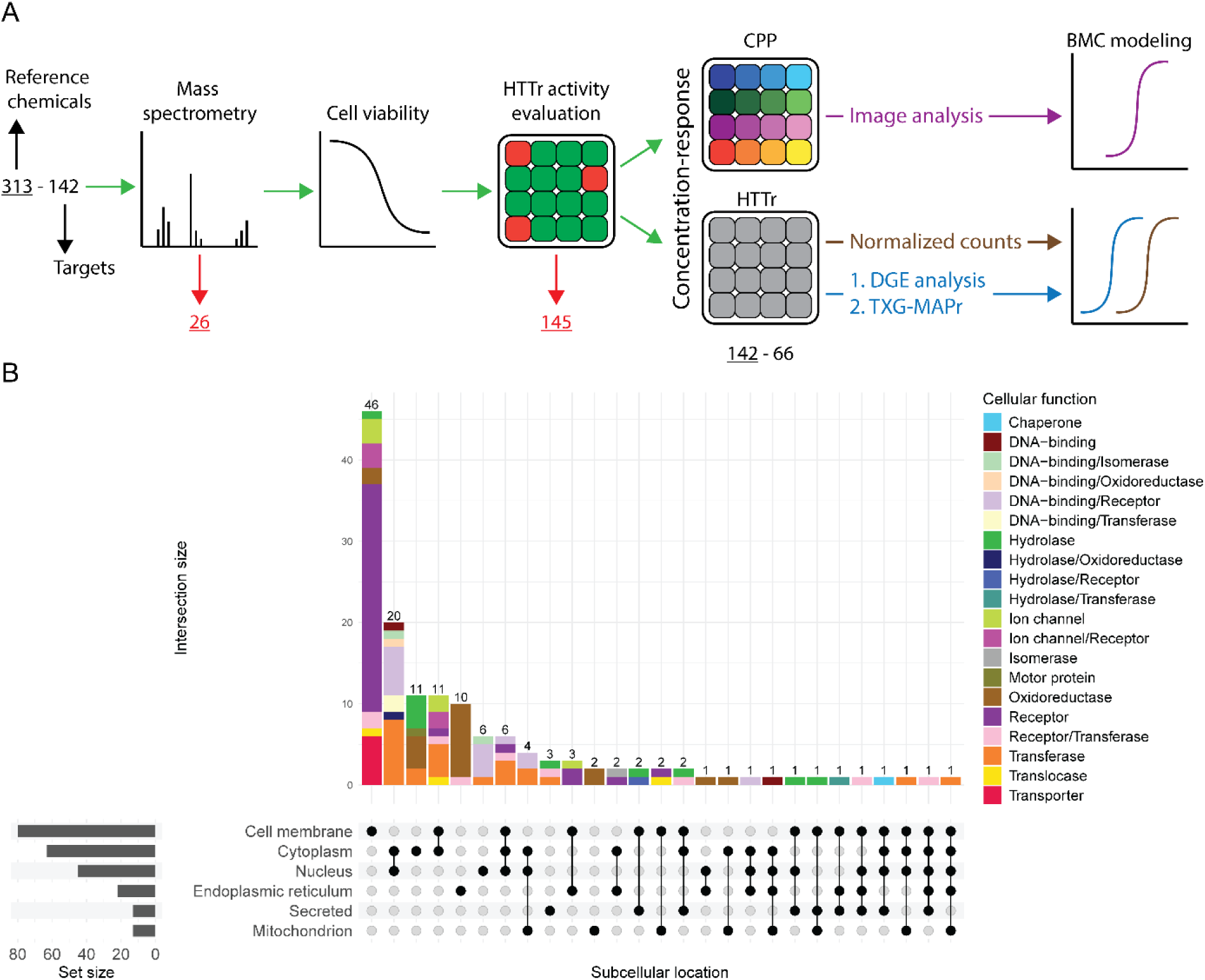
Experimental workflow and UniProt-based biological annotation of reference-chemical target proteins. **(A)** Chronological overview of the experimental workflow, presented from left to right. A set of 313 reference chemicals spanning 142 targets was assembled and subjected to identity quality control by mass spectrometry. Chemicals that could not be identified were excluded (n = 26). A cell viability assay was used to determine the highest non-cytotoxic concentration for each chemical in RPTEC/TERT1. HTTr activity was then assessed at this concentration, and compounds with no detectable response were classified as HTTr-inactive and excluded (n = 145). Concentration-response profiling was subsequently performed across seven concentrations in parallel for CPP and HTTr, testing 142 reference chemicals spanning 66 targets. BMC modeling was performed separately for CPP and HTTr concentration-response data. For CPP, BMC modeling was used to derive morphological activity profiles and phenotype-altering concentrations. For HTTr, BMC modeling was used to determine module tPODs using both transformed normalized count data and TXG-MAPr-derived ME profiles. RPTEC/TERT1 TXG-MAPr modules were used as predefined gene sets for BMC aggregation into module tPODs. **(B)** Functional and subcellular localization annotation of protein targets. Gene symbols of target proteins were mapped to UniProt identifiers and used to query the UniProt API to retrieve cellular function and subcellular location annotations. Retrieved annotations were manually curated to reduce redundancy to enhance visualization.

### 3.1 Reference chemical HTTr and module tPOD activity

We systematically evaluated the HTTr data of the reference chemicals with respect to DEGs and gene module tPODs. Differential gene expression analysis identified active conditions primarily at the upper end of the tested concentration range for each chemical, whereas only a limited number of chemicals exhibited a concentration-dependent increase in the number of differentially expressed genes across the full tested range (Fig. 2, left panel). Sporadic spikes in gene expression responses were observed for several chemicals that lacked consistent activity at the upper end of the concentration range (red labels in Fig. 2 left panel), including Y27632 dihydrochloride (ROCK1 inhibitor), DTXSID6040743 (RARA agonist), everolimus (mTOR inhibitor), cladribine (RRM1 inhibitor), both AVPR2 antagonists and drospirenone (PGR antagonist). For some chemicals, abrupt transitions from inactive to highly active states were observed, potentially indicative of steep potency curves (green labels in Fig. 2 left panel), as seen for colchicine (TUBA1A inhibitor) and alsterpaullone (GSK3B inhibitor). On the gene level, the number of DEGs at the highest concentration and the number of log2(CPM)-derived gene tPODs (in orange in Fig. 2) were generally of the same order of magnitude, indicating concordant activity between the DGE profiling and BMC modeling approaches. In addition, DEG activity at the upper end of the concentration range coincided with tPOD distributions centered near the highest tested concentration. On the module level, a few chemicals (blue labels in Fig. 2) exhibited a very high number of modules with valid module tPODs, approaching the total of 291 modules in the human in vitro RPTEC/TERT1 TXG-MAPr. For instance, DTXSID50648017 (CDK2 inhibitor, 243 modules), silmitasertib (CSNK2A1 inhibitor, 259 modules), 1,1′-Diethyl-2,2′-cyanine iodide (SLC22A1 inhibitor, 224 modules), and alsterpaullone (GSK3B inhibitor, 248 modules). We then compared the two tPOD derivation methods at the module level. Although the number of module eigengenes (ME)-derived module tPODs was consistently lower than that of the log2(CPM)-derived tPODs, their overall distributions were similar. However, in some instances, the ME-derived approach did not identify any valid module tPODs.

**Figure 2.**
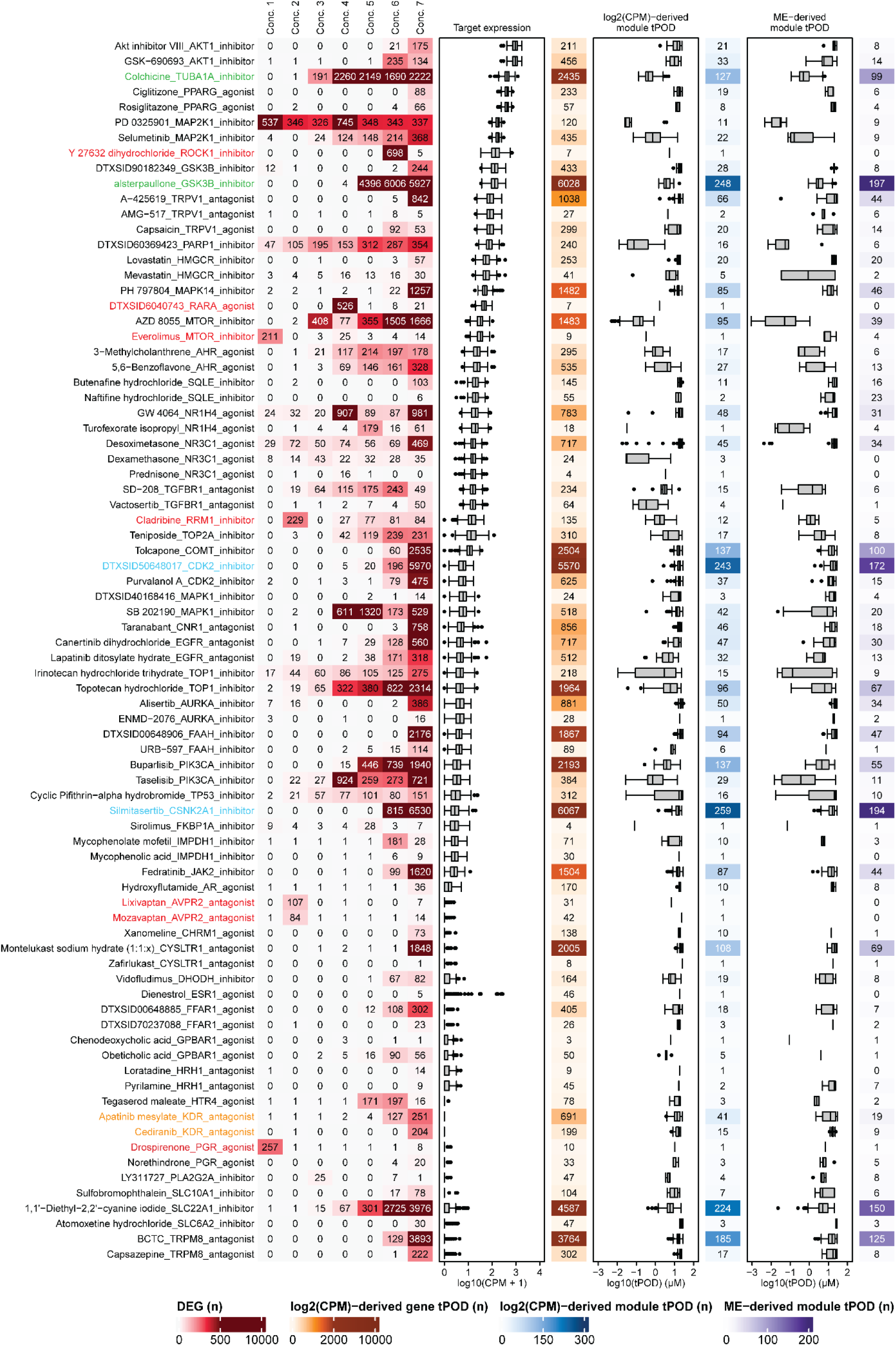
HTTr-active reference chemicals: DGE profiles, target expression, and module tPOD distributions. Each row represents one reference chemical annotated by target and modulation type. Labels shown in color are discussed explicitly in the main text. The left panel shows the number of DEGs across tested concentration range. DEGs were defined as genes with an absolute log2FC > 1.5 and adjusted p-value < 0.05. Concentrations are represented as indexed levels 1 (lowest) to 7 (highest). Target expression indicates baseline normalized expression of the annotated target in RPTEC/TERT1 quantified across 217 negative control samples. Reference chemicals targets are ordered by their median normalized expression value. The number of valid log2(CPM)-derived gene tPODs is indicated in orange. Module tPOD distributions are shown for log2(CPM)-derived and TXG-MAPr ME-derived module tPODs. Values in blue and purple in the adjacent columns indicate the number of modules with valid tPODs for each approach, respectively. Chemicals were classified as HTTr-active if gene set aggregation of log2(CPM)-derived gene tPODs identified at least one module.

The 142 chemicals that were tested in a concentration response were classified as HTTr-active (n = 80) when gene set aggregation of log2(CPM)-derived gene tPODs yielded at least one module, and as HTTr-inactive (n = 62) otherwise. Among HTTr-active (Fig. 2) and -inactive chemicals (Fig. S3), the normalized baseline expression levels of the targets in the negative controls showed a general correlation with DEG activity. HTTr-active chemicals with low baseline target expression levels were generally less active than those with high baseline target expression levels, relative to the overall distribution of target expression levels. Surprisingly, given that KDR (VEGFR) (orange labels in Fig. 2) expression is not expressed (log10(CPM+1) = 0), the observed DEGs and module tPODs possibly reflect off-target transcriptomic responses. This observation suggests that reference chemicals nominally associated with the same target can nonetheless segregate into HTTr-active and HTTr-inactive groups, plausibly reflecting compound-specific ADME properties or limited molecular-level physiology to emulate these interactions within the investigated RPTEC/TERT1 cell system. Consistent with this notion was the observation that the 62 chemicals in the HTTr-negative group were generally related to very low expressed targets (Fig. S3). Among these targets were serotonin receptors HTR2A, HTR1D, and HTR4, which are normally expressed in neuronal synapses.

### 3.2 TXG-MAPr ME-based experimental correlations

We anticipated strong similarity in transcriptomic activity between conditions within and across reference chemical target pairs in the HTTr-active group. To assess this, we included all individual compound and concentration conditions with at least 20 DEGs. For these selected conditions, full transcriptomics profiles were uploaded to the human in vitro RPTEC/TERT1 TXG-MAPr platform to generate quantitative gene network ME activity profiles, which were subsequently used for correlation analysis (Kunnen et al., 2024; van Kessel et al., 2026). Several cellular targets were represented by only a single chemical with only one active condition, limiting the reliability of interpretation between conditions and chemicals. Consequently, further analysis was restricted to pairs of active chemicals acting on the same cellular target (Fig. 3). Similarly, a correlation analysis based on DGE profiles was performed, revealing patterns consistent with those from the ME-based analysis (Fig. S4). High within-chemical condition correlations were observed for several compounds, including 5,6-Benzoflavone (AHR agonist), lapatinib (EGFR antagonist), AZD 8055 (MTOR inhibitor), purvalanol A (CDK2 inhibitor), and DTXSID50648017 (CDK2 inhibitor), despite the limited number of tested conditions (two per compound). DTXSID90182349 (GSK3B inhibitor) also exhibited strong correlations, likely reflecting widespread transcriptional perturbation consistent with high numbers of DEGs. PD 0325901 and selumetinib (MAP2K1 inhibitors) showed consistent and strong within-chemical correlations across multiple active conditions. Taselisib and buparlisib (PIK3CA inhibitors) as well as topotecan and irinotecan (TOP1 inhibitors) showed strong within-chemical correlations. Notably, TOP1 inhibitors exhibited the highest concordance at the highest tested concentrations. Inter-chemical correlations across targets revealed moderate similarity among compounds acting within related signaling pathways. PIK3CA, AKT1 and MTOR inhibitors showed cross-target correlations, suggesting coordinated modulation of the PI3K/AKT/mTOR signaling cascade and downstream transcriptional responses. The strength of these correlations varied among compounds, reflecting differences in target selectivity and the extent of downstream pathway inhibition. Similarly, EGFR antagonists and MAP2K1 inhibitors exhibited notable inter-target correlations, indicating convergent transcriptional effects consistent with their roles within the MAPK signaling cascade. These results suggest that compounds targeting sequential nodes within major signaling pathways elicit concordant transcriptomic outcomes despite differences in their primary cellular targets.

**Figure 3.**
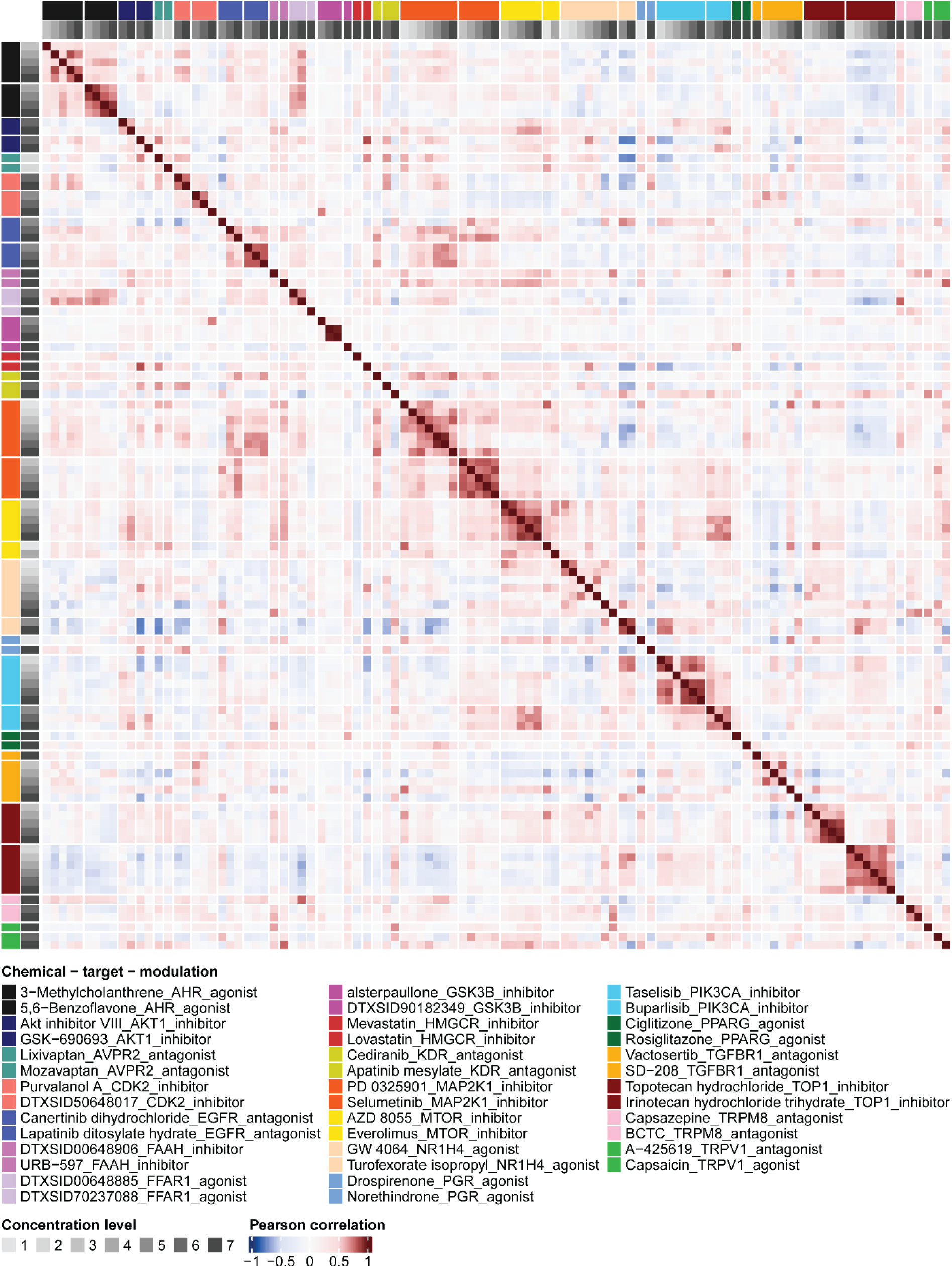
RPTEC/TERT1 TXG-MAPr ME-based biological grouping of HTTr-active reference chemical pairs. Reference chemicals are grouped by target and displayed in the same order on both the x- and y-axis, as defined by the chemical-target-modulation legend. Axis ordering and color annotations follow the legend order. The legend is arranged in three columns and should be read top-to-bottom within each column, proceeding from the left column to the right column. Concentrations are represented as indexed levels 1 (lowest) to 7 (highest). HTTr-active conditions were selected for ME-based correlation analysis if they contained at least 20 DEGs. The Pearson correlation color scale was adjusted to emphasize correlation coefficients greater than 0.5 and smaller than −0.5.

### 3.3 Module-based biological grouping captures downstream effects of MAP2K1 and EGFR inhibition

We further substantiated the mechanisms underlying the inter-target transcriptional correlations in the context of the gene module biology. First, transcriptional responses to MAP2K1 inhibitors and EGFR antagonists were examined by integrating DGE analysis, TXG-MAPr module correlation patterns, and BMC modeling to elucidate shared and opposing pathway activities. Selumetinib (MAP2K1 inhibitor) and both EGFR inhibitors (lapatinib and canertinib) exhibited clear DGE concentration–response profiles (Fig. 4A). PD 325901 (MAP2K1 inhibitor) induces consistent transcriptional perturbation, with sporadic peaks in upregulated genes observed at 0.03 and 0.95 µM. Mechanistic similarity between EGFR and MAP2K1 inhibition was examined using the RPTEC/TERT1 TXG-MAPr by performing correlation analysis at the highest tested concentration, where the DGE response was most pronounced. Biological grouping based on ME correlation analysis with a module activity threshold of |ME| > 2 revealed that MAP2K1 inhibitors and EGFR antagonists consistently downregulated modules 12, 88, 95, and 260 (Fig. 4B). Module annotation and over-representation analysis indicated that modules 88 and 95 were associated with MAPK signaling, while modules 12 and 260 were linked to cell cycle regulation (Tab. S2). Consistent with the expected direction of pathway perturbation, PD 0325901 (MAP2K1 inhibitor) showed moderate negative correlation (r = −0.45) with hEGF (EGFR ligand), with modules 88 and 95 oppositely perturbed. Across all ME correlations, module 95 showed the strongest response, and examination of its gene members through differential expression analysis confirmed that the log2FC magnitude aligned with the corresponding ME trends (Fig. 4C). For reference, our previous module 95 log2FC data from the hEGF treatment derived at 4- and 8 h were included to illustrate the opposing transcriptional response (van Kessel et al., 2026). BMC modeling resulted in varying number of valid log2 CPM-derived gene tPODs for module 95 (Fig. 4D). Confidence intervals were narrow, and the module tPOD (vertical dotted line, median gene tPOD) overlapped with most individual gene BMC confidence intervals. For module 95, module tPODs obtained using the ME-based approach were of the same order of magnitude as those derived from the log2(CPM)-based method. The ME-derived tPODs for modules 12, 88, 95, and 260 were consistent with the overall trend in chemical potency, reflecting both the onset and magnitude of differential gene expression within module 95. Collectively, TXG-MAPr-based biological grouping and BMC modeling indicated that MAP2K1 inhibitors and EGFR antagonists elicit largely convergent MAPK transcriptional signatures, with both classes producing responses opposite to hEGF stimulation and MAP2K1 inhibitors exhibiting greater potency.

**Figure 4.**
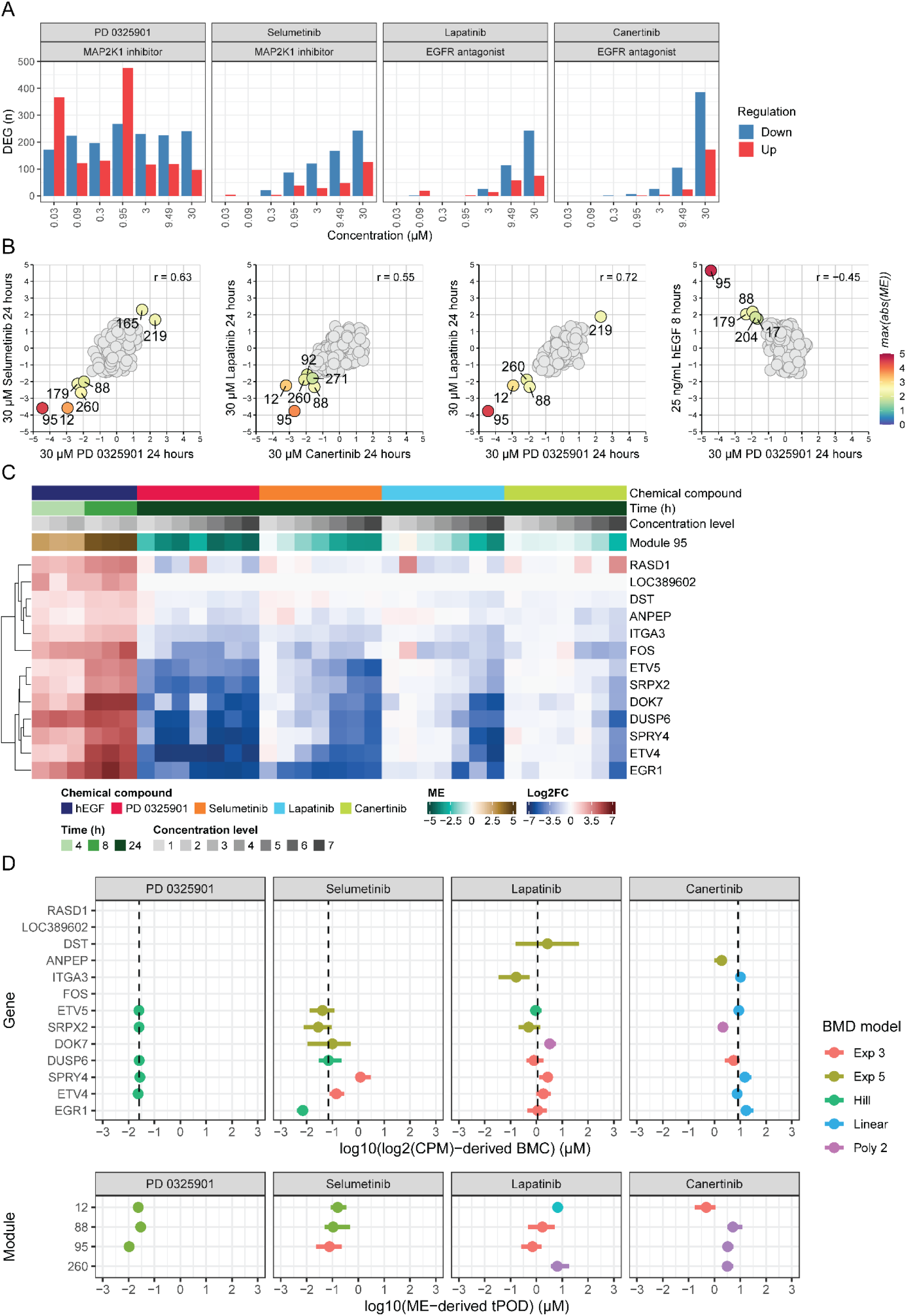
RPTEC/TERT1 TXG-MAPr-based inference of mechanism of action for EGFR antagonists and MAP2K1 inhibitors. **(A)** Differential gene expression profiles and **(B)** TXG-MAPr-based biological grouping of EGFR antagonists and MAP2K1 inhibitors at the highest tested non-cytotoxic concentration at 24 h. DEGs were defined as genes with log2FC > 1.5 (upregulated) or log2FC < −1.5 (downregulated) and adjusted p-value < 0.05. Biological grouping indicated moderate similarity across treatments driven by concordant activity, exemplified by EGFR signaling modules 88 and 95 and a cell-cycle module 12. For every pairwise comparison, the annotations indicate the three most strongly perturbed modules for positive and negative modulation. Annotations were merged when the same module was identified more than once. **(C)** Module 95 ME and log2FC activity profiles. hEGF was included to highlight the opposite response pattern relative to EGFR pathway inhibition. **(D)** The top panel reports gene-level BMC confidence intervals (BMCL-BMC-BMCU) for module 95 gene members where a valid BMC estimate was obtained. The dotted line indicates the median gene-level BMC, which represents the log2(CPM)-derived module tPOD. The bottom panel shows ME-derived module tPOD confidence intervals for modules 12, 88, 95, and 260.

Closer examination of the ME-based correlations revealed that the highest concentration of cediranib, a KDR inhibitor, showed strong correlation to the response profiles of MAP2K1 inhibitors and EGFR antagonists (Fig. 5A). A similar module activation pattern was observed between PD 0325901 and cediranib (r = 0.77), and lapatinib and cediranib (r = 0.67), with modules 12, 88, and 95 consistently downregulated in both comparisons (Fig. 5B). pChEMBL values indicated secondary activity of cediranib on the EGFR receptor, which was not found for apatinib mesylate. Both cediranib and apatinib mesylate demonstrated activity in the nanomolar range for the VEGFR2 (KDR) receptor, whereas only cediranib showed additional activity toward EGFR, also within the nanomolar range (Fig. 5C). A complete overview of the retrieved pChEMBL values for the KDR inhibitors is presented in Figure S5. Highlighted in Figure 5D are the modules for which a valid module tPOD was determined and shared between the comparisons shown in Figure 5B. Module tPODs were consistent between the log2(CPM)-derived and ME-based approaches, except for modules 204 and 219. As observed previously, clear differences in chemical potency were evident, with the KDR inhibitor requiring the lowest concentration to inhibit the EGFR signaling pathway. In conclusion, the ME-based inter-target correlation matches both pathway-related target as well as off-target-related effects and are reflective of biological responses anticipated from the target inhibition.

**Figure 5.**
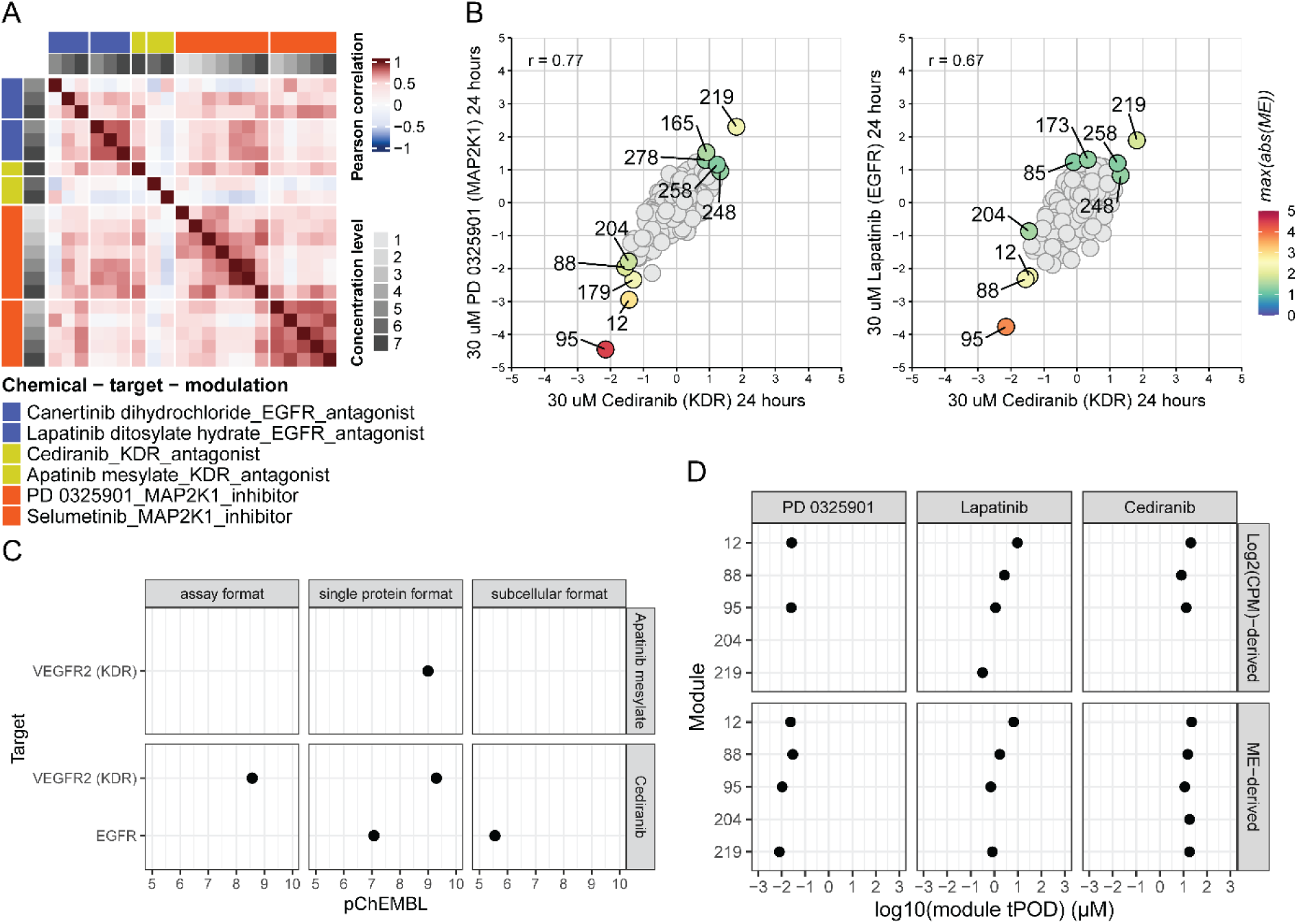
Downregulation of EGFR signaling by off-target activity of KDR antagonist Cediranib. **(A)** RPTEC/TERT1 TXG-MAPr ME-based biological grouping heatmap, shown as a subset of Figure 3 restricted to EGFR and KDR antagonists and MAP2K1 inhibitors. **(B)** RPTEC/TERT1 TXG-MAPr ME-based grouping of PD 0325901 (MAP2K1 inhibitor) and lapatinib (EGFR antagonist) compared with cediranib (KDR inhibitor) at the highest tested non-cytotoxic concentration at 24 h. The three most strongly perturbed modules for positive and negative modulation are annotated and were merged when the same module was identified more than once. **(C)** pChEMBL values for VEGFR (KDR) and EGFR for both KDR inhibitors. pChEMBL values were annotated by assay origin, with assay format, single protein format, and subcellular format indicating the experimental origin from which each pChEMBL value was obtained. **(D)** Log2(CPM)- and ME-derived module tPODs for modules overlapping among the three most perturbed modules (up- or down-modulated) across the three reference chemicals included in the two comparisons shown in panel B. Modules ranked among the top three but not shown in this panel did not yield valid tPODs.

### 3.4 Modulation of cholesterol biosynthesis by reference chemicals

ME-based correlation analysis also demonstrated an expected mechanistically similar activity among PIK3CA, AKT1 and MTOR inhibitors (Fig. 3); these targets act in the PIK3CA/AKT/MTOR pathway. This motivated direct comparison of the inhibitor concentration response characteristics using DGE profiles, TXG-MAPr-based biological grouping and assessment of module tPODs. Taselisib and buparlisib (PIK3CA inhibitors) demonstrated a concentration-related response, while GSK690693 and Akt inhibitor VIII (AKT1 inhibitor) elicited activity only at the upper end of the tested concentration range. Everolimus (MTOR inhibitor) did not show graded activity as concentration increased, whereas AZD 8055 (MTOR inhibitor) exhibited a robust concentration-dependent effect (Fig. 6A). Biological grouping showed that AZD 8055 (MTOR inhibitor) had strong correlation with buparlisib (PIK3CA inhibitor; r = 0.77), followed by Akt inhibitor VIII (AKT1 inhibitor; r = 0.66), and taselisib (PIK3CA inhibitor; r = 0.40) (Fig. 6B). Across inhibitor classes, a subset of modules repeatedly occupied the top three up- and down-regulated positions, with module 68 being consistently represented and showing activity of |ME| > 2. Surprisingly, module 68 was upregulated by Akt inhibitor VIII (AKT1 inhibitor), while being downregulated by taselisib and buparlisib (PIK3CA inhibitors) and AZD 8055 (MTOR inhibitor). Only module 68 exhibited opposite behavior in activity between inhibitor classes and was therefore the primary focus of subsequent analysis. Over-representation analysis showed that module 68 was annotated for the cholesterol biosynthesis pathway (WP197; coverage 12/15, padj = 5,8E-30) (Tab. S3). In addition, module transcription factor enrichment showed significant overlap with regulons of sterol regulatory element binding protein 1 and 2 (SREBF1/SREBF2). Their regulons significantly intersected with module 68, encompassing FDFT1, HMGCR, HMGCS1, INSIG1, SCD, SQLE (p-value = 1.27e-14), and FDFT1, HMGCR, INSIG1, SQLE (p-value = 6.07e-10), respectively (Tab. S2). Interestingly, HMGCR was among the gene members of module 68 and part of the cholesterol biosynthesis gene set. Inspection of activity scores across all chemicals with |ME| > 2 showed that HMGCR inhibitors mevastatin and lovastatin upregulated module 68 (Fig. 6C). Although module 68 was strongly activated by both HMGCR inhibitors, no correlation was detected between them (r = −0.02) (Fig. 6D). Consistent with prior results, module tPOD estimates from log2(CPM)- and ME-derived approaches were of similar magnitude (Fig. 6C). Module 68 tPOD could not be resolved for turofexorate isopropyl (NR1H4 agonist) by either method. For taselisib (PIK3CA inhibitor), a tPOD for module 68 was obtained exclusively with the ME-based method. Module 68 received the top tPOD rank for both HMGCR and PIK3CA inhibitors using the ME-derived tPOD approach. In addition, both inhibitor classes exhibited a broad distribution of module tPODs across the tested concentration range, except for lovastatin which produced module tPODs between 10 and 30 µM (Fig. 6E). Notably, module 68 ranked lowest for Akt inhibitor VIII (AKT1 inhibitor) by both tPOD determination methods. The tPODs for AZD 8055 (MTOR inhibitor) were confined to a lower tPOD distribution range, which reflected the highest tested concentration of 1.5 µM. Collectively, these analyses identify module 68 as a cholesterol biosynthesis signature that is differentially modulated across the PI3K/AKT/mTOR axis, with PIK3CA and MTOR inhibition predominantly suppressing this pathway and HMGCR inhibition inducing compensatory upregulation.

**Figure 6.**
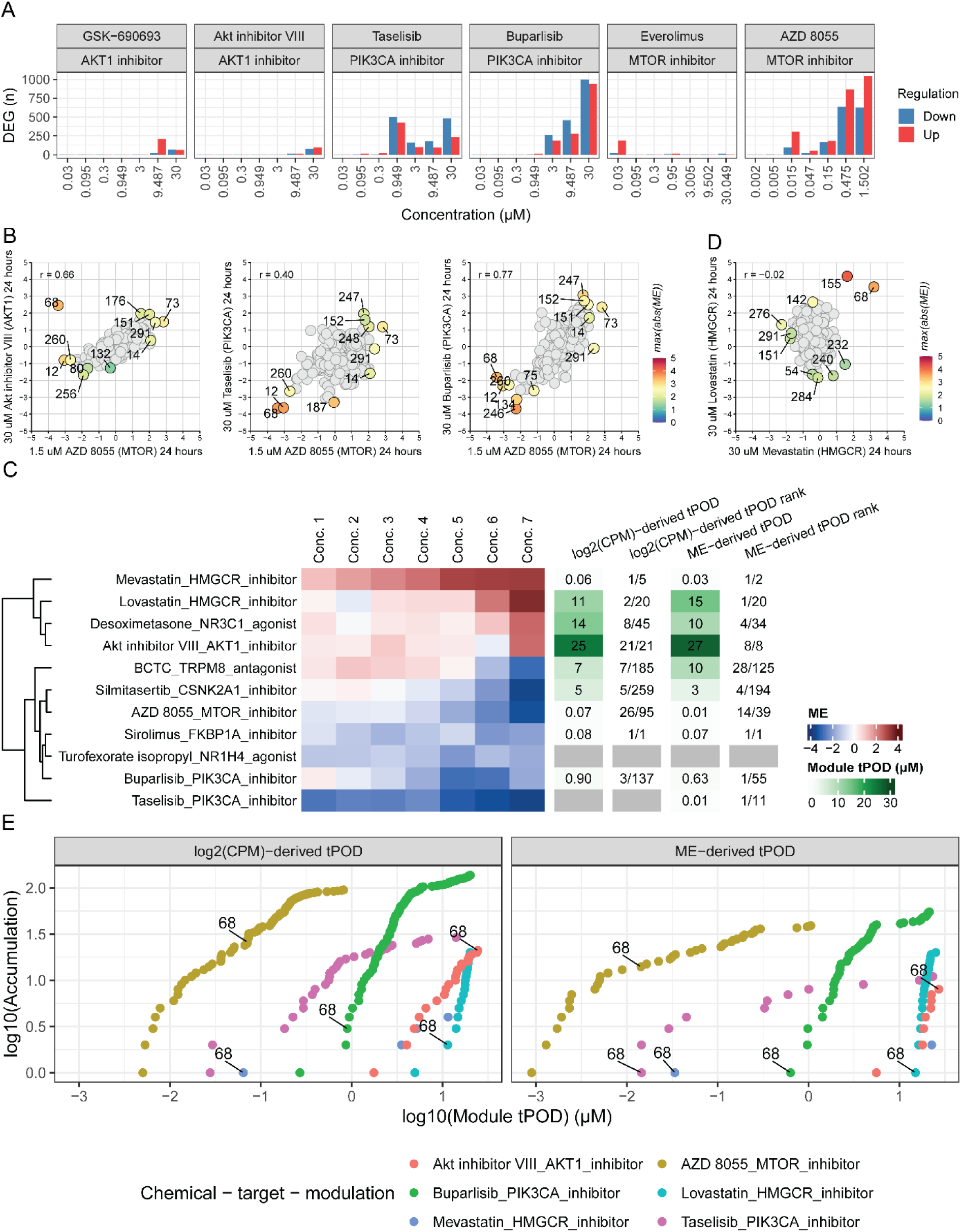
Targeted modulation of the PIK3CA–AKT1–mTOR axis drives divergent cholesterol metabolic responses in RPTEC/TERT1 cells. **(A)** Differential gene expression profiles of PIK3CA, AKT1 and MTOR inhibitors. DEGs were defined as genes with log2FC > 1.5 (upregulated) or log2FC < −1.5 (downregulated) and adjusted p-value < 0.05. **(B)** RPTEC/TERT1 TXG-MAPr ME-based grouping of Akt inhibitor VIII (AKT1 inhibitor) and the two PIK3CA inhibitors, compared with AZD 8055 (MTOR inhibitor), at the highest tested non-cytotoxic concentration at 24 h. The three most strongly perturbed modules for positive and negative modulation are annotated and were merged when the same module was identified more than once. **(C)** The left panel shows the ME profile of module 68 for reference chemicals with ME > 2 in at least one condition, clustered using complete-linkage hierarchical clustering. Concentrations are represented as indexed levels 1 (lowest) to 7 (highest). The right panel shows log2(CPM)- and ME-derived tPODs for module 68, with the accumulation rank shown in adjacent columns. **(D)** RPTEC/TERT1 TXG-MAPr ME-based grouping of HMGCR inhibitors. Module annotation is identical to that described in panel B. **(E)** BMC accumulation profiles for PIK3CA, AKT1, MTOR, and HMGCR inhibitors, with module 68 highlighted in curves for which a module tPOD was available.

### 3.5 Cell Painting Plus: HTTr tPOD vs CPP PAC

To assess the correlation between transcriptomic and phenotypic responses, we performed phenotypic profiling of the reference chemicals using the CPP assay. First, CPP-active reference chemicals were defined as compounds exhibiting at least one active feature category, where an active feature category was defined as containing at least 30% of features with significant BMCs. These chemicals were then compared with the hits identified in the transcriptomic screen. All HTTr-active reference chemicals exhibited at least one feature category (channel-region) with at least 30% significant features, except for PD 0325901 (MAP2K1 inhibitor) (Fig. 7A), which showed detectable activity but did not reach the threshold (Fig. 7A). By comparison, HTTr-negative reference chemicals showed reduced CPP activity across feature categories, with 50 chemicals exceeding the CPP activity threshold and 12 remaining CPP-inactive (Fig. S6). Broad activity across multiple feature categories was exemplified by silmitasertib (CSNK2A1 inhibitor) (orange label in Fig. 7A) and extended across several additional HTTr-active reference chemicals spanning diverse target classes. To illustrate the extent of morphological perturbation under these conditions, representative images are provided in Figure S7. Beyond this highly active group, several chemicals exhibited more selective morphological responses. Clustering of feature category activities revealed enriched responses in lysosome-channel cytoplasm and ring features for everolimus (mTOR inhibitor), mycophenolate mofetil (IMPDH1 inhibitor), both PGR agonists, and the two transporter inhibitors atomoxetine hydrochloride (SLC6A2 inhibitor) and sulfobromosphthalein (SLC10A1 inhibitor) (green labels in Fig. 7A). In contrast, DTXSID40168416 (MAPK1 inhibitor) and sirolimus (FKBP1A inhibitor) (purple labels in Fig. 7A) predominantly activated the mitochondria-channel cytoplasm and ring features. Notably, although everolimus predominantly affected lysosome-related features and sirolimus predominantly affected mitochondrial features, both compounds also induced weaker effects in the reciprocal feature category. This overlap is consistent with their shared downstream inhibition of mTOR signaling. However, the differences in their dominant phenotypic responses may reflect distinct proximal mechanisms of action, as everolimus directly targets mTOR, whereas sirolimus exerts its effects through binding to FKBP1A, which subsequently inhibits mTOR (Hu et al., 2021; Witzig et al., 2015). Additionally, the two AVPR2 inhibitors showed strong responses in both lysosome and mitochondria-related feature categories and were clustered together (brown labels in Fig. 7A).

**Figure 7.**
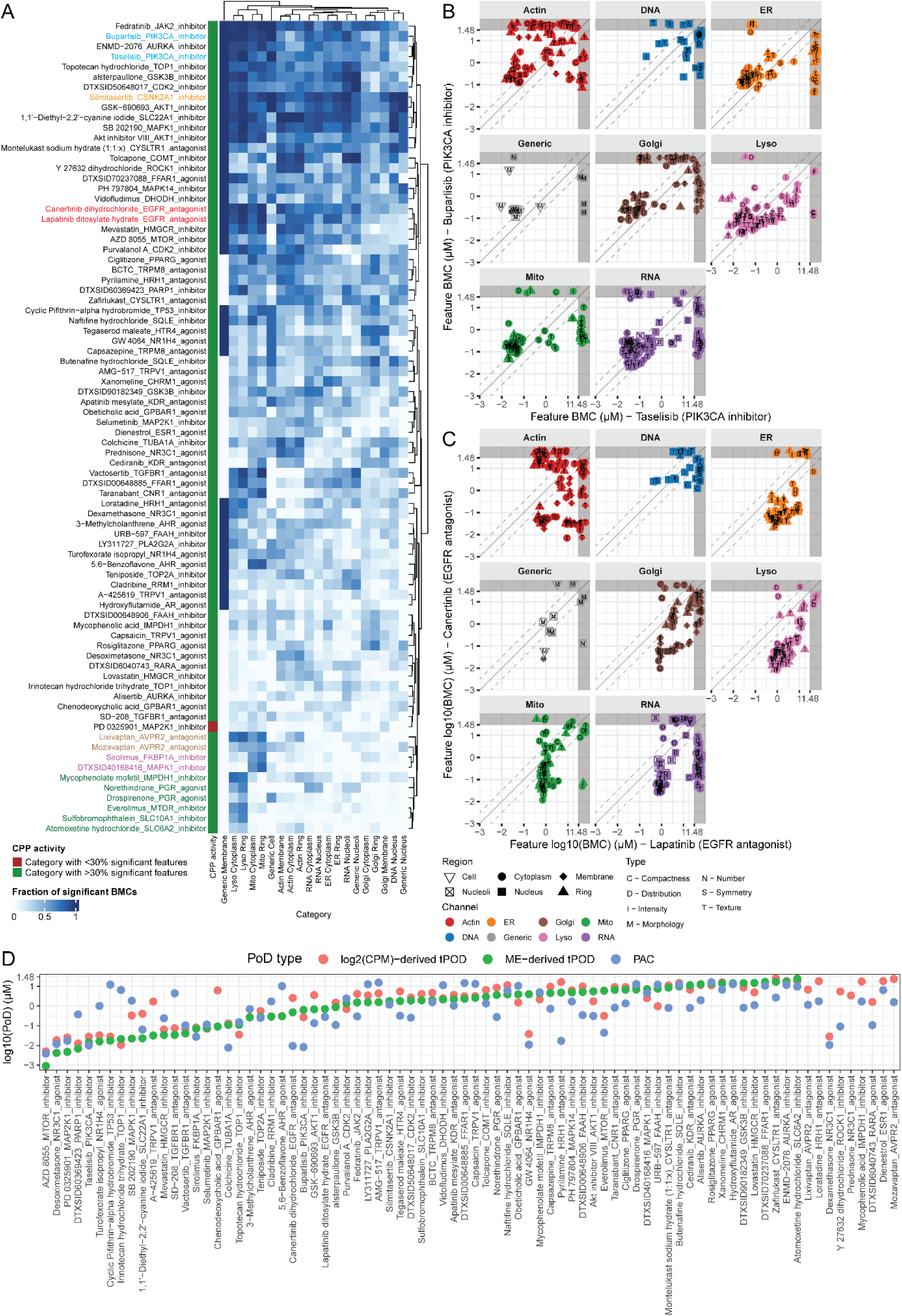
CPP activity among HTTr-active reference chemicals. **(A)** The fraction of significant feature BMCs per feature category defined by imaging channel and cellular region is reported for each reference chemical annotated by target and modulation type. Reference chemicals were classified as CPP-active when at least one feature category contained more than 30% significant features. Labels shown in color are discussed explicitly in the main text. **(B)** CPP feature BMCs for PIK3CA inhibitors and **(C)** CPP feature BMCs for EGFR antagonists. The solid line indicates the 1:1 relationship and the dotted lines indicate a threefold difference. Grey bands beyond 30 µM (log10 = 1.48) denote the plotting region for incomplete feature BMC pairs, where a valid feature BMC was obtained for only one of the two reference chemicals. **(D)** Comparison of CPP- and HTTr-derived points of departure for HTTr-active reference chemicals. For each chemical, the lowest log2(CPM)-derived module tPOD, the lowest ME-derived module tPOD, and the CPP-derived PAC are reported. Reference chemicals are ordered by increasing ME-derived module tPODs. The two module tPOD minima are derived independently within each method and therefore may correspond to different modules.

Among the chemicals with high activity across feature categories, PIK3CA and AKT1 inhibitors were in proximity within the clustering (blue labels in Fig. 7A), in accordance with the HTTr data. Detailed analysis of feature BMCs revealed highly similar response patterns for both PIK3CA inhibitors, with buparlisib exhibiting slightly lower potency across actin, endoplasmic reticulum, generic, Golgi apparatus and mitochondria channels (Fig. 7B). In contrast, buparlisib elicited a stronger transcriptional response, perturbing more modules with tPODs across a broad range (Fig. S8A). In addition, clustering of feature category activities showed high similarity between EGFR inhibitors (red labels in Fig. 7A). Canertinib consistently exhibited greater potency across features of the actin, endoplasmic reticulum, Golgi apparatus, lysosomes, mitochondria, and RNA channels (Fig. 7C). Although both antagonists perturbed actin-related features, no clear relationship was discernible. For canertinib, the module tPOD distribution was centered between 10 and 30 µM (Fig. S8B) whereas CPP feature BMCs were generally lower, indicating higher apparent phenotypic potency relative to the transcriptomic response.

Next, PACs from CPP were summarized for each compound using previously published method based on Mahalanobis distance (Nyffeler et al., 2020) for comparison of compound potency estimates between HTTr and CPP. Among the available comparisons, 23 of 77 log2(CPM)-derived tPODs (29.9%) and 23 of 69 ME-derived tPODs (33.3%) were within one log10 unit of the corresponding CPP-derived PAC, representing a tenfold range. Thus, approximately one-third of the tPOD estimates from each workflow were within tenfold of PAC, whereas most tPOD estimates differed from PAC by more than one order of magnitude. These descriptive proportions were based on different numbers of available comparisons (Fig. 7D). The compounds showing close alignment in ME-derived tPOD included several kinase inhibitors targeting the PI3K/AKT/mTOR and MAPK signaling pathways, such as taselisib (PIK3CA inhibitor), sirolimus (FKBP1A/mTOR pathway modulation), AZD8055 (mTOR inhibitor), ENMD-2076 (AURKA inhibitor), and multiple MAPK inhibitors. Growth factor receptor inhibitors also exhibited similar concordance, including vactosertib (TGFBR1 inhibitor) and lapatinib (EGFR inhibitor). However, compounds targeting the same protein did not consistently exhibit similar tPOD–PAC relationships, indicating that concordance was driven more by chemical-specific response patterns than by shared nominal targets. Overall, concordance between module tPODs and CPP-derived PACs was chemical-specific rather than systematic, with no consistent relationship between transcriptomic and phenotypic potency across the HTTr-active reference set. Taken together, both transcriptomic and phenotypic data captured coherent responses for chemicals affecting core kinase signaling pathways, particularly PI3K/AKT/mTOR and EGF/MAPK. While HTTr and CPP can act complementary for hazard identification, HTTr will provide mechanistic underpinning of target-related responses that support mode-of-action similarity.

## 4 Discussion

In this study, we established paired concentration-response datasets in RPTEC/TERT1 cells, comprising HTTr and CPP profiles for 142 mass spectrometry-verified reference chemicals that affect 66 different targets, tested across seven concentrations at 24 hours. HTTr responses of chemical-target pairs strongly correlate but require concentration response datasets that are reflective of different target engagement potencies. HTTr response similarities are observed for targets that act in the same signaling pathway and allow to verify possible off-targets effects. In general, CPP supports hazard identification for targets that are highly expressed. Both HTTr and CPP biological perturbations are mainly observed for targets with sufficient and detectable baseline gene expression. Our data indicate that TXG-MAPr gene module-derived tPOD can contribute to mechanism-based hazard assessment, in support with CPP-derived PAC evaluation. We processed the HTTr data using our previously developed RPTEC/TERT1 TXG-MAPr network (van Kessel et al., 2026). Scores derived from the network modules highlighted transcriptomic similarities among compounds associated with the same target or pathway. Importantly, overlaying new data onto the existing RPTEC/TERT1 network, which was derived from a dataset comprising stress-response compounds and nephrotoxicants, enabled mechanistically relevant interpretation. This finding demonstrates the breadth of mechanisms captured by the initial network, which resulted from careful compound selection. Further refinement of the RPTEC/TERT1 network could involve constructing an updated version using a curated subset of the HTTr data obtained in this study.

The reference chemicals used in this study aim to modulate specific cellular targets and are therefore expected to elicit comparatively focused transcriptional responses (Judson et al., 2018b), manifested as coherent activation or repression of a limited set of pathway-associated modules. High HTTr correlations indicate broadly concordant modulation across many modules, which was consistent with a pervasive and uniform transcriptional response. However, in this context, low to modest correlations may reflect focused, pathway-specific responses in which mechanistically relevant activity was confined to a small set of strongly responsive modules (van Kessel et al., 2026). Under these conditions, target engagement can remain evident at the level of individual module responses. This was exemplified by cholesterol biosynthesis module 68 following exposure to HMGCR inhibitors, even when correlation across the full module space is low. For future applications of the TXG-MAPr framework, such as chemical read-across and grouping, these findings emphasize the need for well-curated reference datasets that capture both broad transcriptional similarity and focused, pathway-specific module responses. Such datasets would reduce the risk of overlooking mechanistically related chemicals solely because they exhibit low global HTTr correlations.

As a result of enzymatic inhibition of HMGCR, cholesterol biosynthesis module 68 was strongly upregulated at 24 hours. This response is consistent with SREBF1- and SREBF2-regulated transcriptional feedback following pathway inhibition (Horton et al., 2002). Notably, the direction of module 68 regulation differed across inhibitors of the PI3K/AKT/mTOR axis (DeBose-Boyd & Ye, 2018). PIK3CA inhibitors and the mTOR inhibitor AZD 8055 reduced cholesterol biosynthesis whereas AKT inhibitor VIII induced activity. The ability to detect and distinguish opposing regulatory directions at the module level adds interpretive value for triage and follow-up design, even when causal resolution is beyond the scope of the present HTTr analysis. In general, we provide a characterization of targets that can be reasonably perturbed and whose responses can be quantified in RPTEC/TERT1 cells, thereby adding an additional layer of evidence for the use of TXG-MAPr-based chemical grouping for NGRA purposes.

For the HTTr chemical-target correlation, we used a minimum threshold of 20 DEGs. This criterion was implemented as a practical activity filter to prioritize chemical-target pairs for concentration-response comparisons. However, some chemicals may induce highly specific target-mediated responses that affect only a limited number of sentinel genes, particularly at low or moderate concentrations. Such responses may have been excluded despite evidence of target engagement. In contrast, chemicals satisfying the 20-DEG criterion may partly reflect broader compensatory, adaptive, or stress-related transcriptional changes. Consequently, this filtering step may have favored chemicals with more extensive transcriptomic perturbations and may have underrepresented narrow, highly specific chemical-target signatures. This criterion relates to a broader extent on how to define a so called “active” chemicals, whose selection might impact downstream analysis such as correlation analysis or BMC modelling (Costa et al., 2024).

Following this concept, methods for deriving tPOD can strongly impact the assessment of chemical potency and NAM method sensitivity (O’brien et al., 2025). In this study, we elaborated on two approaches for computing TXG-MAPr-based module-level tPODs, namely log2(CPM)- and ME-based and applied additional filtering criteria in both methods (Kunnen et al., 2024). For log2(CPM)-derived module tPODs, each module was required to contain at least one gene identified as a DEG; for ME-derived module tPODs, we retained only modules with an absolute ME greater than 1 in at least one condition for each reference chemical. Across reference chemicals, log2(CPM)- and ME-derived module tPODs were highly correlated, indicating broad consistency between tPOD estimates. For instance, EGFR signaling module 95 and cholesterol synthesis module 68 showed comparable tPODs between methods and were detected across a broad concentration range. However, the primary systematic difference was qualitative rather than quantitative. For a subset of modules, a tPOD was obtained with one method but no tPOD could be derived using the other. These method-dependent dropouts and inclusions limit direct module-level comparability and indicate differential sensitivity to effect magnitude, concentration-response relationships, and signal variability. To date, no study has directly compared tPOD estimates obtained using two distinct workflows: gene-level concentration–response modeling of log2(CPM) data followed by calculation of the median BMC among genes belonging to each module, and direct concentration–response modeling of module eigengene scores. A related direct gene-set approach calculates normalized enrichment scores using single-sample gene-set enrichment analysis, followed by BMC modeling (Harrill et al., 2024). The present study therefore extends the current methodological framework by directly comparing post hoc gene-set summarization of gene-level BMCs with direct modeling of gene-set-level expression scores. Universally optimal BMC configurations and filtering criteria are difficult to justify because platform-dependent measurement properties, dataset characteristics and quality, and preprocessing and filtering decisions jointly determine signal retention and, consequently, BMC-derived potency; therefore, further investigation is required to establish consensus pipelines, including through coordinated initiatives such as PARC projects (Systems Toxicology) and the Health and Environmental Sciences Institute (HESI) Molecular Point of Departure Working Group (O’Brien et al., 2025).

Relative to HTTr, CPP often demonstrated higher apparent sensitivity compared with HTTr and classified a larger proportion of reference chemicals as active. For some compounds, CPP activity profiles were broadly distributed across multiple feature categories, indicating diffuse perturbations affecting numerous cellular features. This combination of a high proportion of active features and widespread perturbations across different channels suggests a broad morphological response but limits mechanistic interpretability. In contrast, other compounds showed more selective engagement of specific phenotypic domains, exemplified by compounds predominantly affecting lysosome and mitochondrial related morphological features. In principle, mechanistic inference from CPP can be strengthened through large-scale guilt-by-association analyses that compare query profiles against well-annotated reference profiles (Ewald et al., 2026). However, within the present experimental design, which included generally two compounds sharing the same target and narrowly defined target classes based on combined protein targets and modes of action (antagonists and agonists), the CPP assay provided less mechanistic insight than transcriptomic profiling. Furthermore, the morphological information was restricted to seven organelles, limiting the depth of biological interpretation. Therefore, CPP activity in this study is best interpreted as evidence of cellular perturbation rather than as a standalone basis for mechanism assignment without additional reference data and comparative profiling. Future studies should substantially expand the number and diversity of chemicals analyzed by CPP in RPTEC/TERT1 cells. A larger drug target specific reference dataset would facilitate robust guilt-by-association analyses and improve the mechanistic interpretation of CPP-derived phenotypic profiles in parallel with HTTr data. Integration of HTTr and CPP data into a shared foundational model could provide complementary mechanistic interpretation and predictive capabilities (Ha et al., 2025; Lejal et al., 2025; Way et al., 2022).

The utility and interpretability of both HTTr and CPP are dependent on context. Activity profiles were generated in RPTEC/TERT1 using reference chemicals with defined cellular targets at a single time point. Accordingly, the performance characteristics and interpretive trade-offs reported here reflect this specific test system, experimental design, and analysis workflow. Direct extrapolation to other in vitro systems is not warranted because assay responsiveness and the distribution of elicited biological effects can differ across models. For instance, intracellular exposure was not explicitly characterized. Proximal tubule transporter activity can influence cellular uptake and efflux and, therefore, the effective intracellular concentration (Aschauer et al., 2015). Consequently, in vitro kinetics will determine whether a reference chemical can reach and interact with its intended target. Incorporating transporter-informed exposure predictions, for example by anticipating efflux liability for specific transporter substrates, would improve interpretation of negative findings. Finally, the present conclusions are based on reference chemicals selected for high-confidence target information. Extension to industry-relevant chemicals that likely will not have specific biological targets, will require greater reliance on grouping based on similarity of the biological response. Accordingly, future work should evaluate HTTr and CPP performance across additional in vitro cell models, exposure durations, and chemical classes to enhance adoption in regulatory decision-making.

In conclusion, HTTr and CPP provide complementary early-tier evidence streams for in vitro nephrotoxicity assessment within a weight-of-evidence framework. Within an NGRA workflow, CPP could serve as a sensitive initial triage screen to detect biological perturbations and prioritize chemicals for subsequent HTTr analysis. HTTr could then provide data-driven mechanistic interpretation through the TXG-MAPr framework within a test-system-specific toxicogenomic context. Ultimately, integrating both modalities within a unified predictive model, together with complementary functional readouts, could further enhance the mechanistic understanding and prediction of cellular responses.

## Data availability

The raw and processed data deposited in ENA and BioStudies repositories will be made publicly available upon acceptance of the manuscript for publication in a scientific journal.

## Supporting information

**Figure S1.**
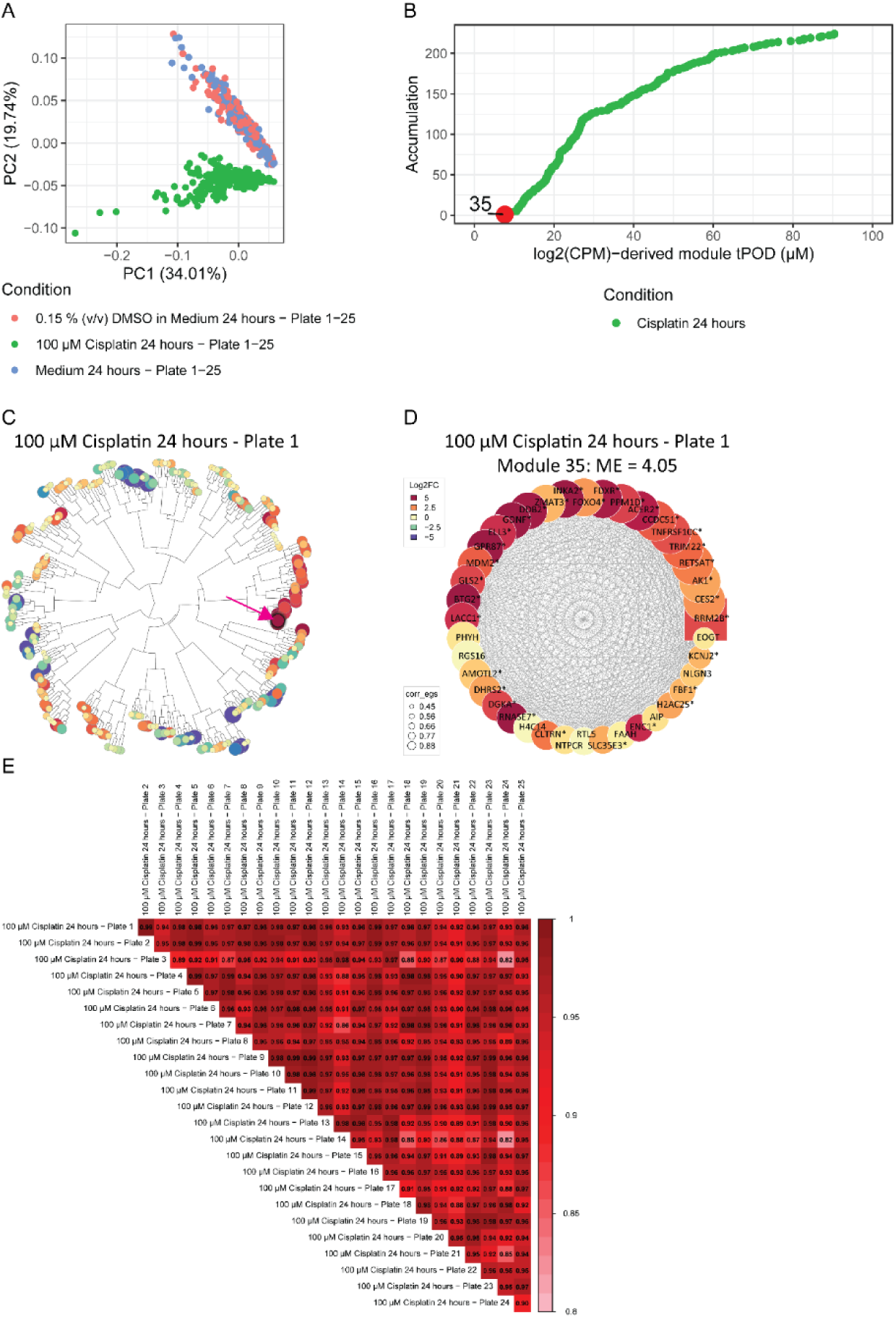
HTTr concentration-response screen quality control. (A) PCA analysis of negative control medium and 0.15% DMSO (v/v) and positive control cisplatin 100 µM at 24 hours after exposure in RPTEC/TERT1. Negative control conditions did not show separation while clear separation between the negative and positive controls was observed. (B) Accumulation of log2(CPM)-derived module tPODs for cisplatin. Module 35 had the lowest accumulation rank. (C) Human in vitro RPTEC/TERT1 TXG-MAPr showing ME projection of 100 µM cisplatin at 24 hours of experimental plate 1. Highlighted is module 35 which showed strong upregulation, indicated by the dark red color. (D) Module 35 gene members shows upregulation which is reflected in the ME score of 4.05. Gene members were associated with the P53-mediated DNA damage response (Tab. S2; Tab. S3). (E) TXG-MAPr-based ME correlation of cisplatin exposure across all experimental plates show high correlation.

**Figure S2.**
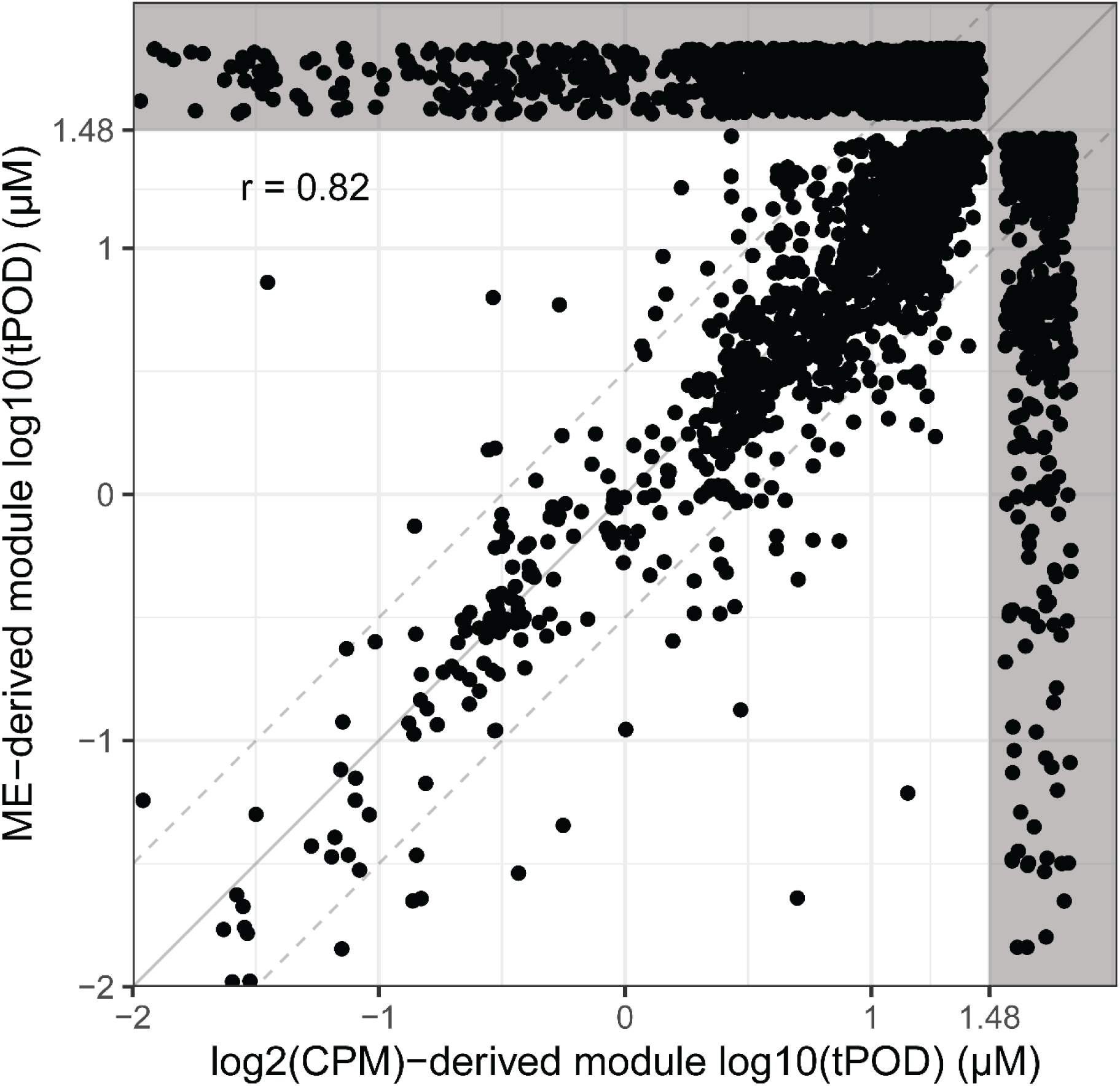
Log2(CPM)- and ME-derived module tPOD correlation. The solid line indicates the 1:1 relationship and the dotted lines indicate a threefold difference. Pearson’s correlation coefficient is reported in the top left corner. Grey bands beyond 30 µM (log10 = 1.48) denote the plotting region for incomplete module tPOD pairs, where a valid module tPOD was obtained through only one of the two methods.

**Figure S3.**
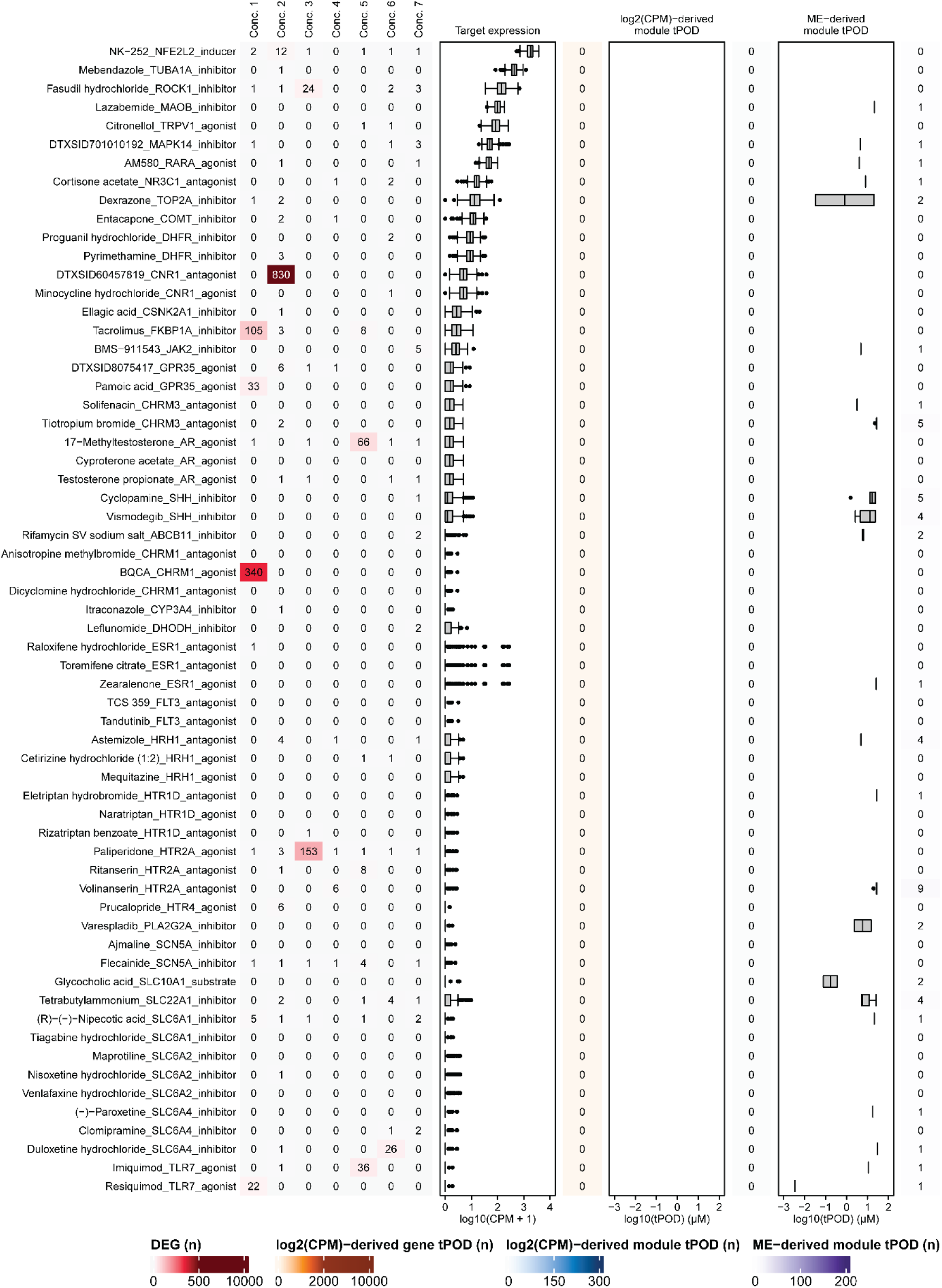
HTTr-inactive reference chemicals. All terms, thresholds, and definitions are as described in the legend of Figure 2.

**Figure S4.**
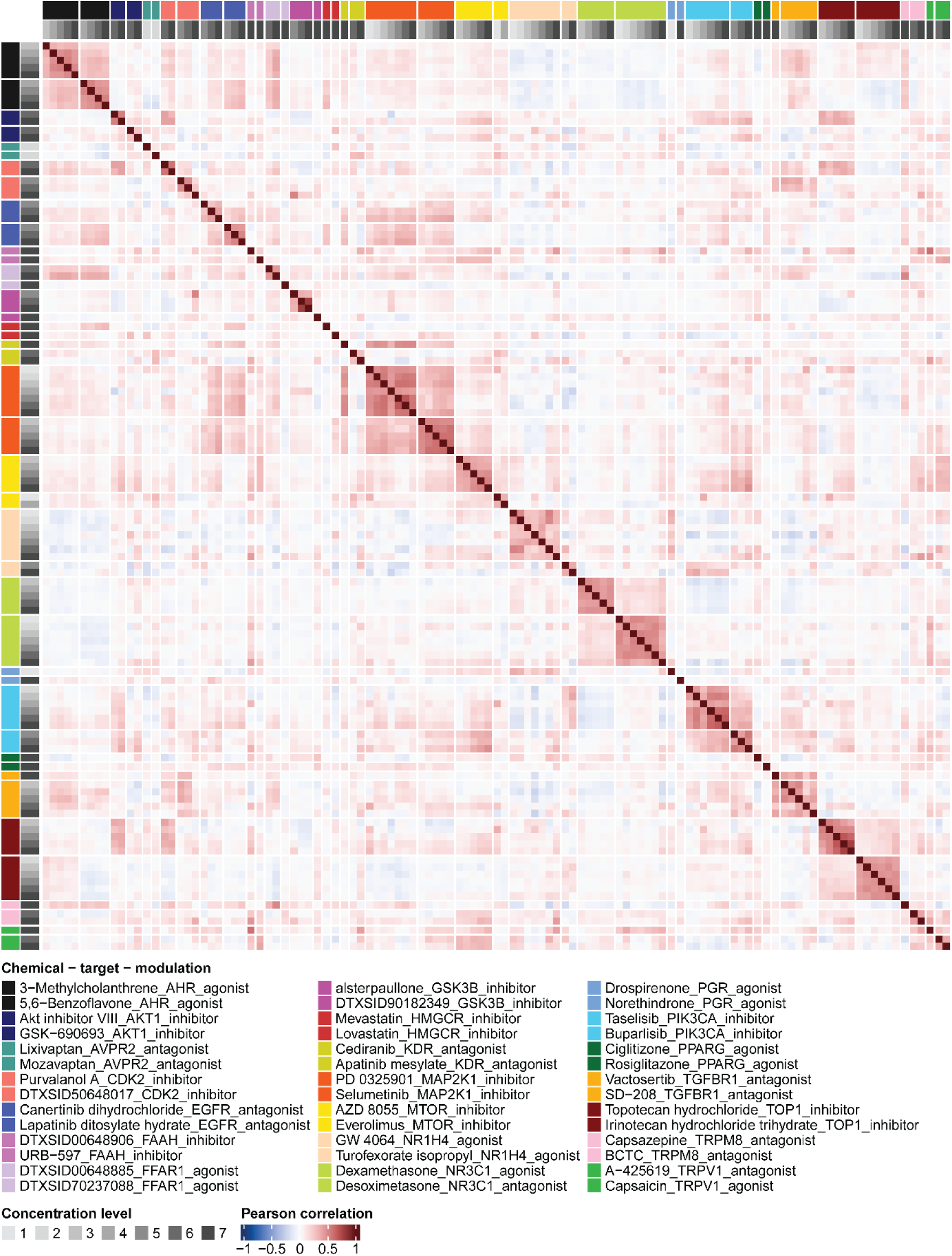
DGE profile correlation of HTTr-active reference chemical pairs. Reference chemicals are grouped by target and displayed in the same order on both the x- and y-axes, as defined by the chemical-target-modulation legend. Axis ordering and color annotations follow this legend order. The legend is arranged in three columns and should be read top-to-bottom within each column, proceeding from the left column to the right column. Concentrations are represented as indexed levels 1 (lowest) to 7 (highest).

**Figure S5.**
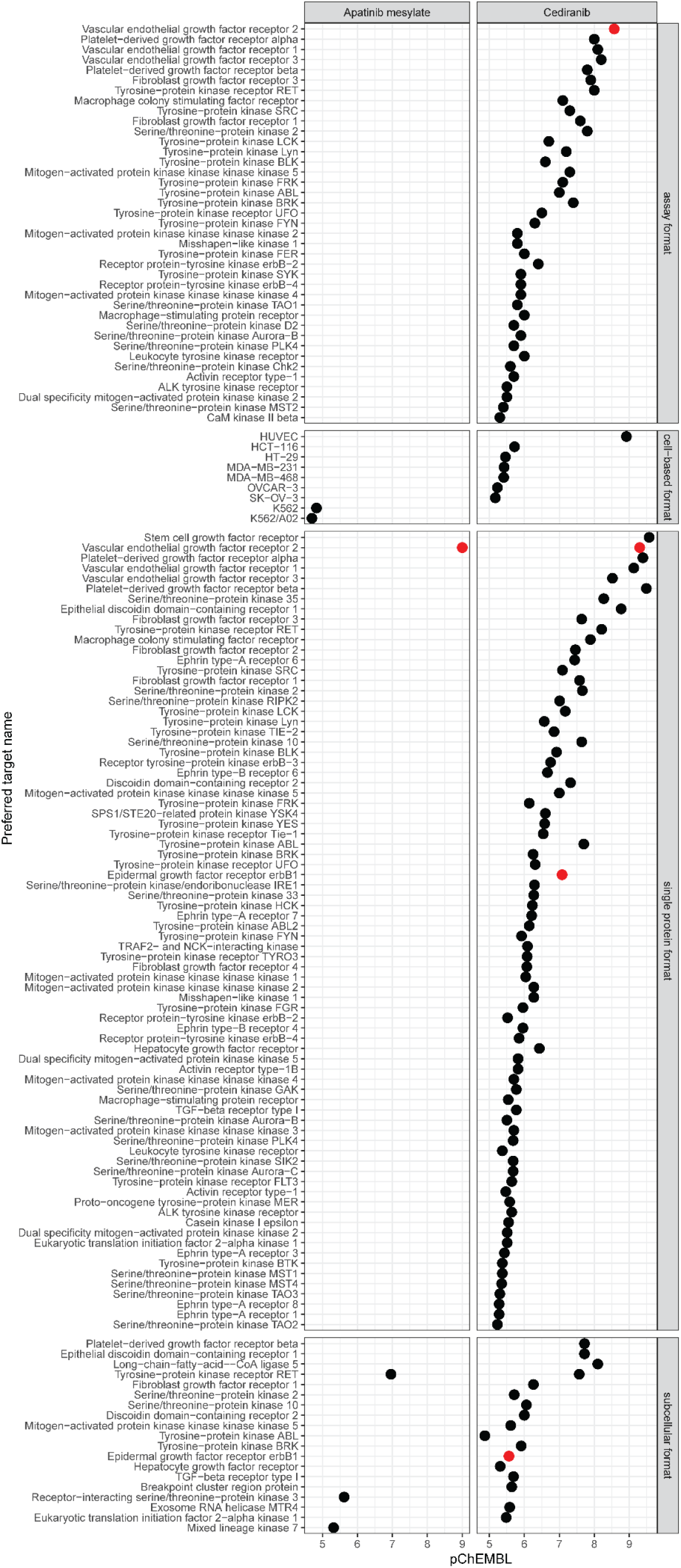
Retrieved pChEMBL values for KDR inhibitors. Highlighted in red are EGFR and VEGFR2 (KDR). The absence of a data point for apatinib mesylate indicates that this compound was not tested in the assay annotated on the right-hand side of the plot.

**Figure S6.**
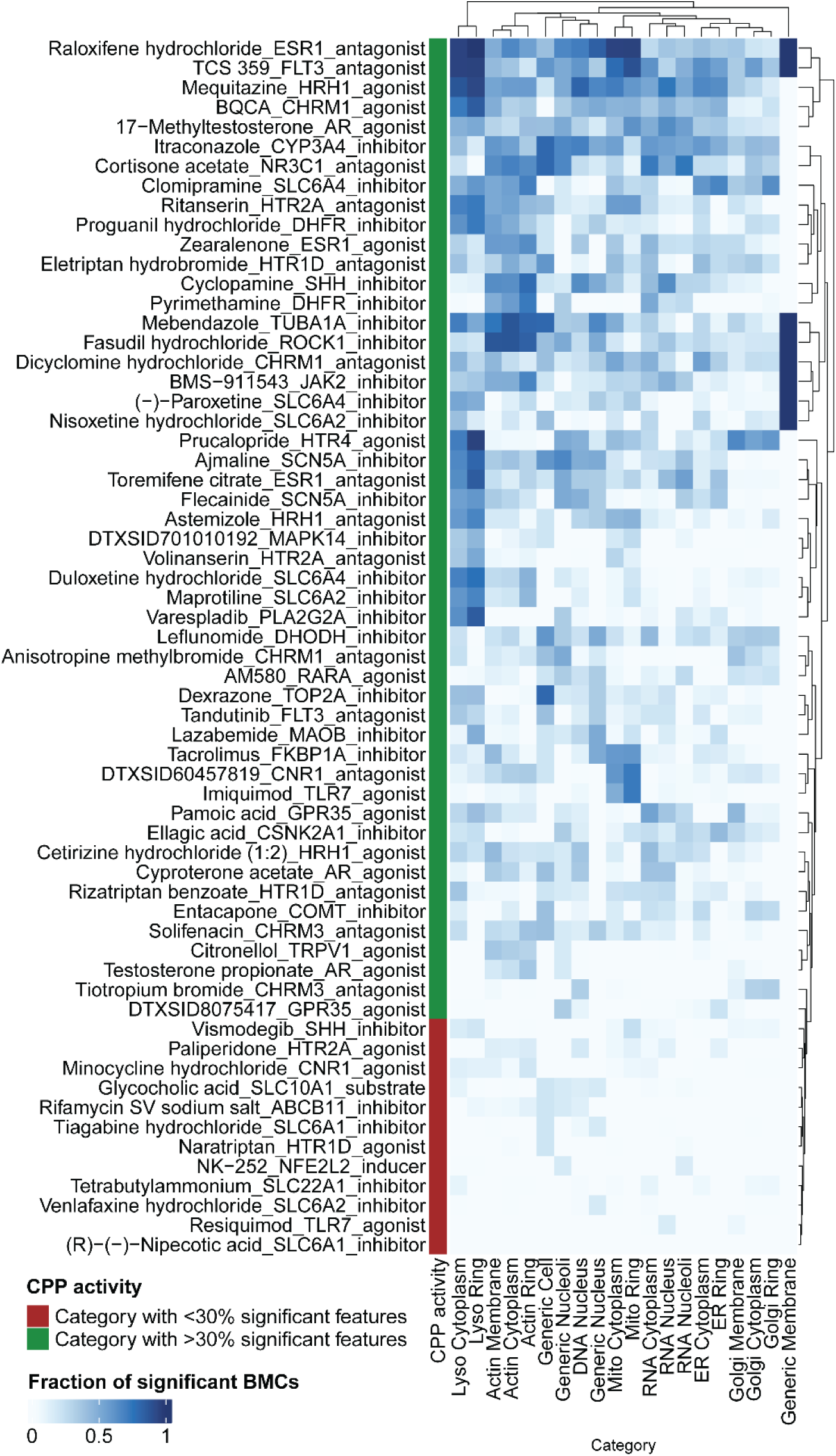
CPP activity among HTTr-inactive reference chemicals. The fraction of significant feature BMCs per feature category defined by imaging channel and cellular region is reported for each reference chemical annotated by target and modulation type. Reference chemicals were classified as CPP-active when at least one feature category contained more than 30% significant features.

**Figure S7.**
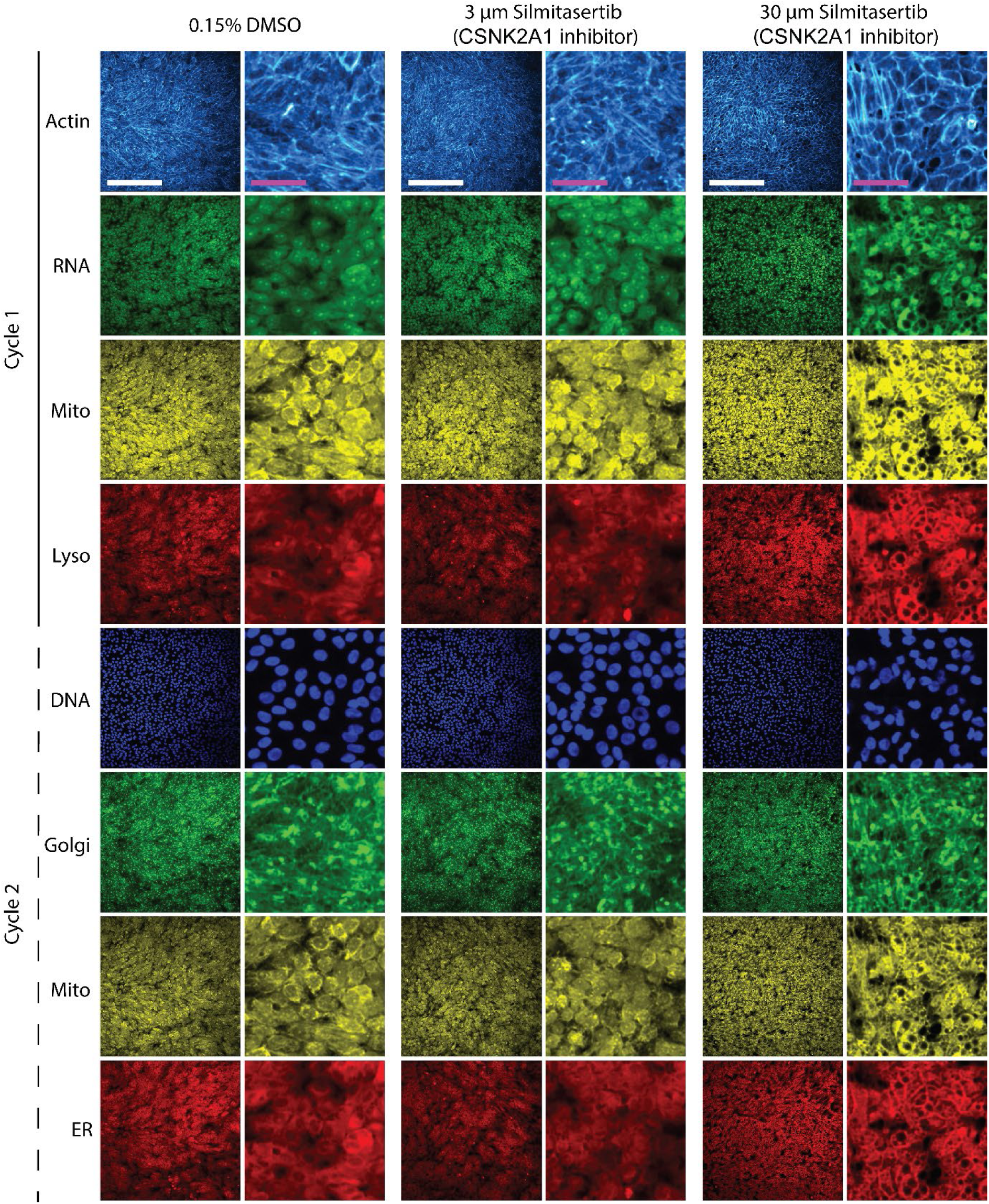
Representative images for silmitasertib (CSNK2A1 inhibitor). Columns (left to right) show 0.15% (v/v) DMSO, 3 µM silmitasertib, and 30 µM silmitasertib. Within each column, the left subpanel shows the full field of view (white scale bar 250 µm), and the right subpanel shows a 5× magnified view (purple scale bar 50 µm). The mitochondrial dye channel was acquired in two imaging cycles. Only the second cycle was used for downstream analysis.

**Figure S8.**
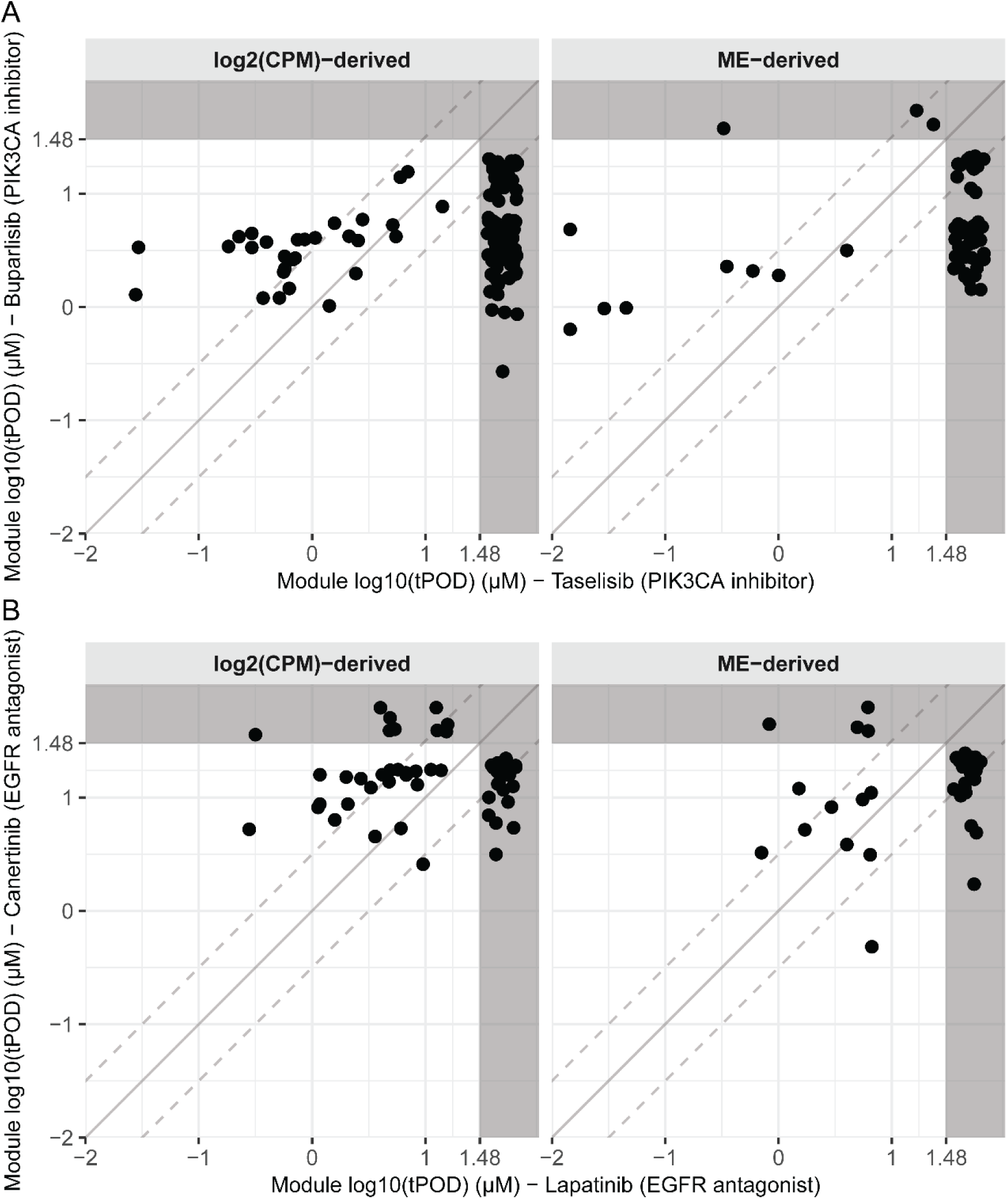
Module tPOD correlation between PIK3CA inhibitors and EGFR antagonists. The solid line indicates the 1:1 relationship and the dotted lines indicate a threefold difference. Grey bands beyond 30 µM (log10 = 1.48) denote the plotting region for incomplete module tPOD pairs, where a valid module tPOD was obtained by only one reference chemical.

